# A distinct chondrogenic progenitor population from human umbilical cord– derived mesenchymal stromal cells for osteoarthritis therapy

**DOI:** 10.64898/2026.11.08.765201

**Authors:** Xiang Gao, Jie Han, Yankui Zhai, Jintong Song, Jia Tian, Lixian Li, Dahong Liang, Zhiping Chen, Jiamei Xu, Yuting Liao, Ahmad, Rui Lin, Liao Cui, Bilian Xu, Li Li, Qixian Xiao, Ren Xu, Yanzhi Liu

## Abstract

Stem cell-based cartilage regeneration remains constrained by functional heterogeneity and phenotypic drift in conventional cell sources, compromising durable structural repair. Human umbilical cord-derived mesenchymal stromal cells (UC-MSCs) represent an accessible, off-the-shelf cell source; however, their broad clinical utility is hampered by cellular heterogeneity, leaving the key chondrogenic subpopulation responsible for cartilage repair unresolved. By integrating single-cell transcriptomics with functional assays, we identified and prospectively isolated a distinct, highly expandable subpopulation of skeletal stem cell-like progenitors (U-CPCs) from the human umbilical cord, establishing an extra-skeletal cell source uniquely primed for joint restoration. Single-cell transcriptomics reveal their inherent regulatory landscape, uncovering a distinct transcriptional state uniquely primed for chondrogenesis. In comparison with conventional stem cells and other tissue-derived SSC-like populations (from umbilical cord, bone marrow, adipose tissue, and dental pulp), U-CPCs exhibit a markedly enhanced chondrogenic capacity. In both a mouse cartilage injury model and a rat osteoarthritis model, the administration of U-CPCs effectively mitigates joint degeneration and drives the restoration of a structurally sound, hyaline-like matrix. Mechanistically, U-CPCs secrete extracellular vesicles enriched with microRNA Let-7a-5p, which specifically modulates DUSP7/MAPK/ERK signaling in recipient chondrocytes, thereby restoring extracellular matrix homeostasis and promoting cartilage repair. By characterizing these specialized extra-skeletal progenitors and elucidating both their intrinsic chondrogenic priming and paracrine regulation of host tissue, this study establishes U-CPCs as a robust, lineage-restricted cellular source for osteoarthritis therapy.

**Graphic Abstract:** 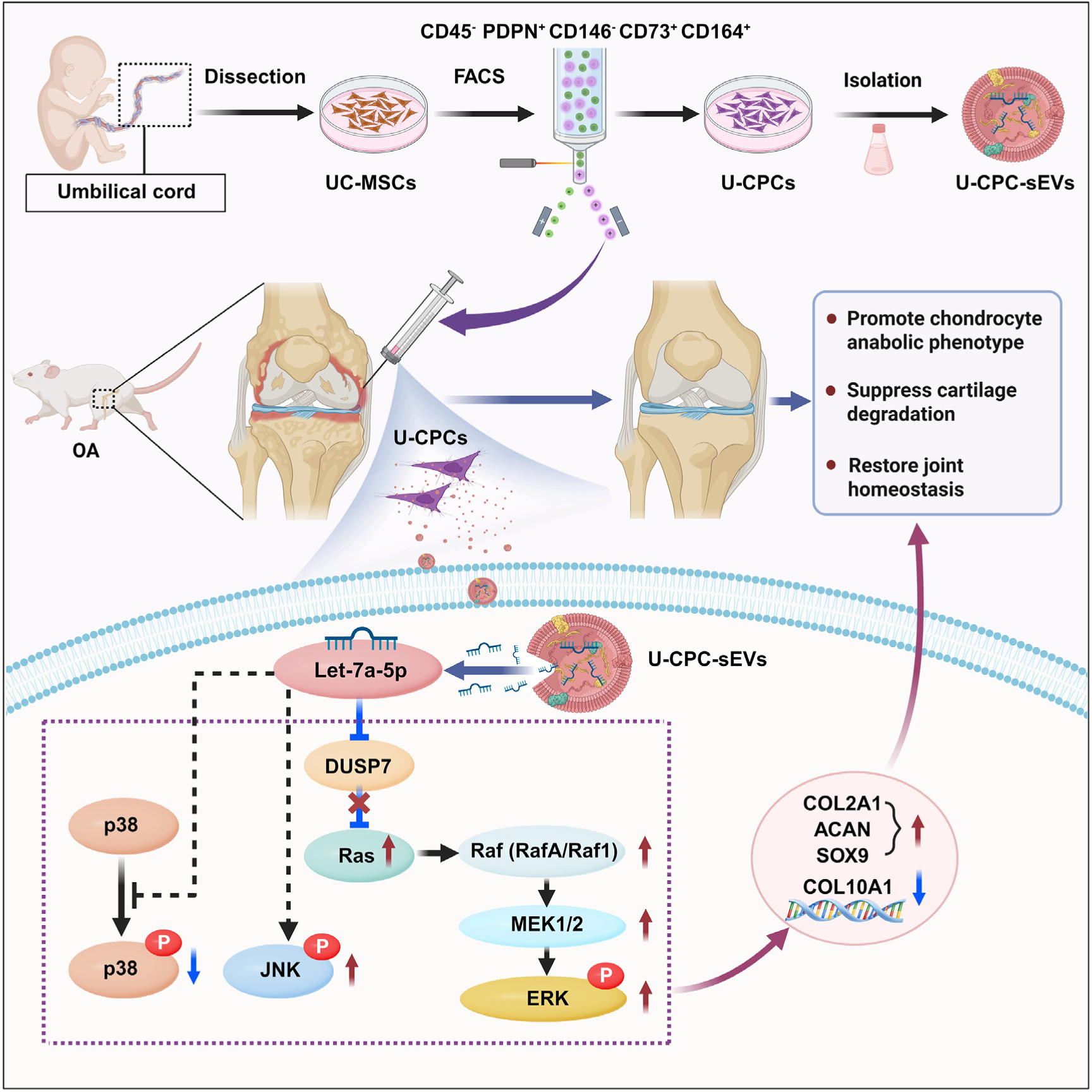

**Highlights:** Identification of lineage-restricted SSC-like progenitors (U-CPCs) in umbilical cord.

U-CPCs maintain stable chondrogenic fidelity without hypertrophy or phenotypic drift.

Intra-articular U-CPCs delivery arrest osteoarthritis and restore hyaline cartilage.

U-CPC-sEVs modulate the let-7a-5p/DUSP7/MAPK axis and reverse cartilage catabolism.

## Introduction

Articular cartilage lacks intrinsic regenerative capacity, making structural reversal of osteoarthritis (OA) a major clinical challenge ^1^. Current interventions, ranging from intra-articular pharmacotherapy to surgical procedures such as microfracture ^2^ and mesenchymal stem cells (MSCs) ^3^ autologous chondrocyte injections ^4^, provide symptomatic relief but fail to restore functional hyaline cartilage, typically resulting in the formation of mechanically inferior fibrocartilage that fails to restore long-term joint homeostasis ^5^. To overcome these limitations, the field is shifting from the use of heterogeneous MSCs to lineage-defined skeletal stem cells (SSCs) ^6^ and skeletal stem and progenitor cells (SSPCs) ^7^. The maintenance of skeletal integrity, which encompasses both chondral repair and bone homeostasis, relies heavily on these specific reservoirs ^8, 9^. Clonal analyses and lineage tracing have molecularly defined murine ^10^ and human SSCs (marked by PDPN^+^CD146^-^CD73^+^CD164^+^) ^11^ distinguishing them from traditional MSCs by their strict epigenetic restriction to osteochondrogenic rather than adipogenic, lineages ^12^.

Recent advances have revealed profound heterogeneity in SSPCs. SSPCs encompass highly specialized, lineage-restricted subsets that occupy distinct postnatal niches, rather than monolithic population ^13^. While various subpopulations have been phenotypically defined (e.g., ZIC1/PAX1- or Procr-expressing cells) ^14, 15, 16, 17, 18, 19, 20, 21^, their collective function remains central to driving postnatal skeletogenesis, fracture regeneration, and joint homeostasis. Despite their regenerative potential, the clinical translation of SSPC-based therapies is limited by two major challenges. First, although recent study has elegantly mapped skeletal stem cell (SSC) hierarchies across various skeletal sites ^22^, the field remains constrained by the strict anatomical localization of these genuine SSCs, low harvest yields from restrictive anatomical niches, and inherent phenotypic instability, notably hypertrophic drift and fibrocartilage deposition. Second, endogenous SSPC pools are highly susceptible to age-related declines. Chronic low-grade inflammation drives both numerical depletion and lineage skewing within the aging niche ^23^, diverting progenitors toward fibroblastic and adipogenic fates at the expense of osteochondrogenesis ^24^. Consequently, relying on endogenous cell recruitment is largely ineffective in older adults with OA. Therefore, establishing a scalable, exogenous source of potent, lineage-committed progenitors that resist senescence-induced phenotypic loss is critical for effective structural joint repair.

Although the human umbilical cord offers an accessible, off-the-shelf cell source of mesenchymal stromal cells (UC-MSCs), the therapeutic efficacy of bulk UC-MSCs remains variable because the specific chondrogenic progenitor subpopulation within this heterogeneous pool is undefined. In this study, we report the prospective isolation and functional validation of a scalable subpopulation of human SSC-like progenitors derived from an extra-skeletal niche: umbilical cord (U-CPCs). Single-cell transcriptomic profiling demonstrates that these U-CPCs possess a primed transcriptional state restricted to the chondrogenic lineage, dictated by a distinct tissue-of-origin epigenetic memory. We demonstrated that U-CPCs possess significantly higher chondrogenic potential than conventional MSCs and homologous SSC-marker-positive populations derived from the bone marrow, adipose tissue, and dental pulp (provisionally designated as B-SSPCs, A-SSPCs, and D-SSPCs, respectively). U-CPCs intrinsically avoids the phenotypic drift typical of adult MSCs, maintaining high-fidelity cartilage deposition. Furthermore, we revealed that U-CPCs drive cartilage regeneration primarily through a paracrine mechanism by secreting small extracellular vesicles (sEVs) that recondition the degenerative chondrogenic microenvironment. Mechanistically, U-CPC-sEVs encapsulate and deliver microRNA let-7a-5p to host chondrocytes, directly modulating the let-7a-5p/DUSP7/MAPK/ERK signaling axis to restore extracellular matrix homeostasis and promote cellular proliferation. Collectively, by characterizing these specialized extra-skeletal progenitors and elucidating both their intrinsic chondrogenic priming and paracrine regulation of host tissue, this study establishes U-CPCs as a scalable, robust, and lineage-restricted cellular source for osteoarthritis therapy.

## Results

### U-CPCs exhibit enhanced proliferation and a chondrogenic-biased differentiation profile

To identify a defined progenitor population with stable chondrogenic potential and improve the consistency and reproducibility of cartilage repair, we prospectively isolated a distinct chondroprogenitor population from UC-MSCs by FACS using a previously validated panel of SSC surface markers ^11^, which we designated as U-CPCs. Quantitative analysis revealed that U-CPCs accounted for approximately 22.7% of the total UC-MSCs population (**Figure 1A**). Following expansion in culture, U-CPCs exhibited the typical spindle-shaped morphology of mesenchymal stem cells (**Figure S2**).

**Figure 1.**
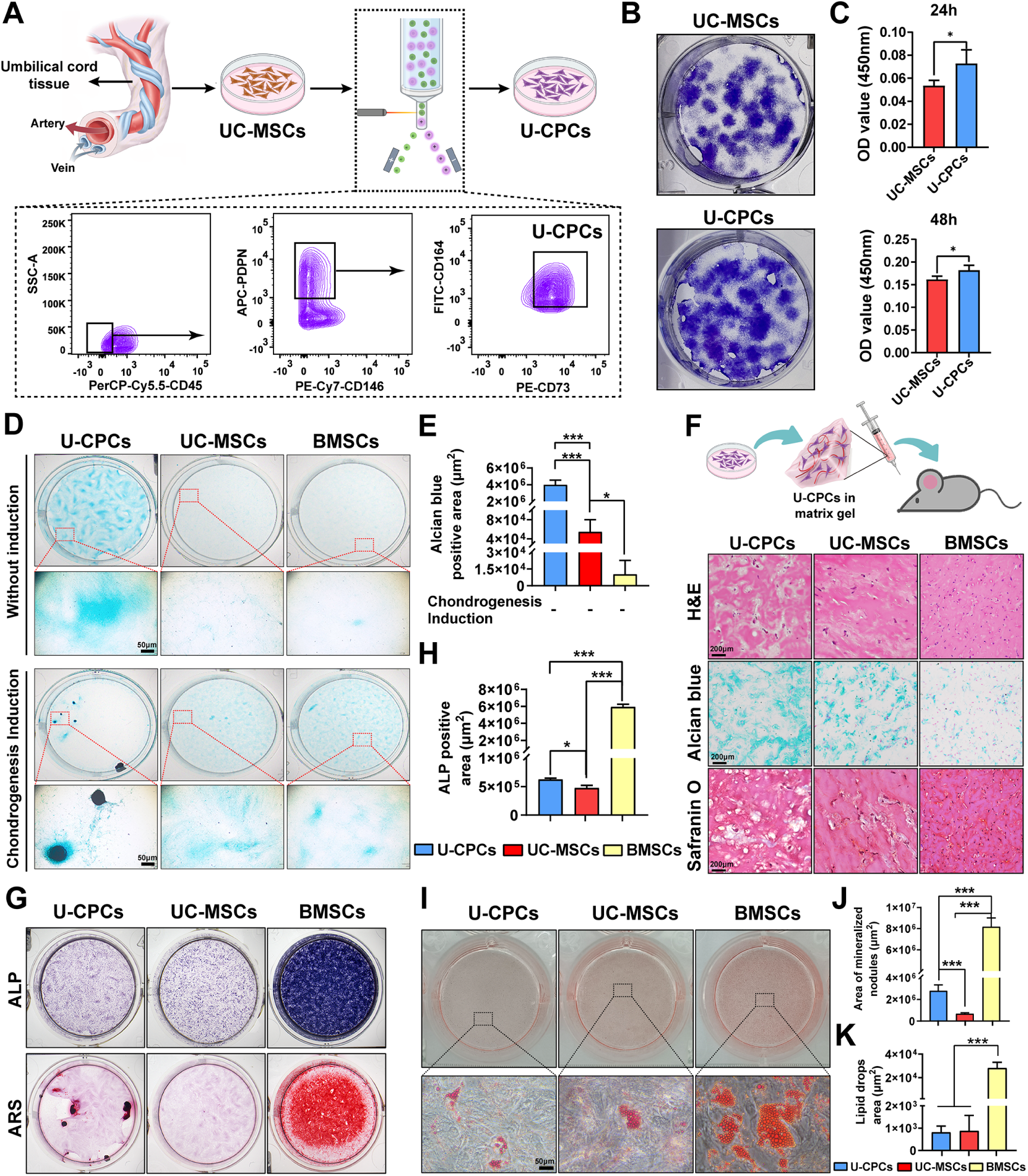
Isolation, viability assessment, and multi-lineage characterization of U-CPCs. (A) FACS isolation of U-CPCs from the UC-MSCs. (B) Comparison of the colony-forming capacity of U-CPCs and UC-MSCs. (C) The viability of U-CPCs and UC-MSCs was evaluated using the CCK-8 assay after 24 or 48 h of culture. (D) Alcian blue staining to assess the chondrogenic differentiation of U-CPCs, UC-MSCs, and BMSCs cultured with or without chondrogenic induction. (E) Quantitative analysis of Alcian blue staining without chondrogenic induction. (F) Histological evaluation of U-CPCs, UC-MSCs, and BMSCs Cell-Matrigel implants after 4 weeks using H&E, Alcian blue, and Safranin O staining. (G) ALP and Alizarin Red S (ARS) staining to evaluate osteogenic differentiation of the indicated cells under osteogenic induction conditions. (H) Quantitative analysis of ALP staining following osteogenic induction. (I) Oil Red O staining for adipogenic differentiation of U-CPCs, UC-MSCs, and BMSCs cultured in adipogenic induction medium. (J) Quantitative analysis of ARS staining after osteogenic induction. (K) Quantitative analysis of Oil Red O staining after adipogenic induction. Data are presented as mean ± SD. \**P*<0.05, \*\**P*<0.01, \*\*\**P*<0.001. Figure 1A was created using BioRender and has confirmed publishing and licensing rights, with agreement number MM2A5EL3VH.

First, we assessed the proliferative properties of U-CPCs using colony formation and CCK-8 assays. After 10 days of culture, U-CPCs displayed a greater colony-forming capacity than UC-MSCs (**Figure 1B**). Consistently, CCK-8 assays demonstrated significantly higher cell viability in U-CPCs than in UC-MSCs at both 24 and 48 h (**Figure 1C**), indicating an enhanced proliferative capacity.

The chondrogenic differentiation potential of U-CPCs was evaluated *in vitro* and *in vivo*. For *in vitro* assessment, U-CPCs, UC-MSCs, and BMSCs were cultured with or without chondrogenic induction medium, followed by Alcian blue staining to assess glycosaminoglycan deposition. Under non-inductive conditions, U-CPCs exhibited greater glycosaminoglycan (GAG) accumulation than UC-MSCs and BMSCs, as evidenced by a larger Alcian blue-positive area. Following chondrogenic induction, U-CPCs formed larger chondrosphere-like structures than the other two cell types (**Figure 1D**), consistent with the quantification of the Alcian blue-positive area (**Figure 1E**). For *in vivo* assessment, each of the three cell types was mixed with Matrigel and subcutaneously implanted into nude mice, and the implants were harvested for histological analysis after four weeks. H&E staining revealed that U-CPC-derived implants exhibited a more compact, cartilage-like architecture than UC-MSCs or BMSCs, while Alcian blue and Safranin O staining further demonstrated greater cartilage matrix deposition in U-CPC-derived implants (**Figure 1F**).

To further define the differentiation profile of U-CPCs, we assessed their osteogenic potential under basal and osteogenic induction conditions. In the absence of induction, U-CPCs and UC-MSCs exhibited weak ALP staining, whereas BMSCs showed relatively higher basal ALP activities (**Figures S3A and S3B**). After 7 days of osteogenic induction, U-CPCs exhibited an intermediate level of ALP activity, which was higher than that of UC-MSCs but lower than that of BMSCs (**Figure 1G**), as further supported by semiquantitative analysis (**Figure 1H**). Mineralization was assessed using Alizarin Red S staining 15 days after induction. Under non-inductive conditions, no appreciable mineralized nodules were observed in any of the three cell types (**Figure S3A**). Following osteogenic induction, U-CPCs formed more mineralized nodules than UC-MSCs but significantly fewer than BMSCs (**Figure 1G**), a pattern confirmed by quantification of the mineralized area (**Figure 1J**). These results indicate that the osteogenic differentiation capacity of U-CPCs is greater than that of UC-MSCs but lower than that of BMSCs.

In contrast to their pronounced chondrogenic potential and intermediate osteogenic capacity, U-CPCs have limited adipogenic differentiation. Under non-inductive conditions, no appreciable intracellular lipid accumulation was observed in U-CPCs, UC-MSCs and BMSCs (**Figure S4**). Following adipogenic induction, Oil Red O staining revealed abundant intracellular lipid droplets in BMSCs, whereas only sparse lipid droplets were observed in UC-MSCs and U-CPCs (**Figure 1I**). Quantitative analysis showed no significant difference in the Oil Red O-positive area between UC-MSCs and U-CPCs, whereas both exhibited significantly less lipid accumulation than BMSCs (**Figure 1K**). Together, these findings establish U-CPCs as a highly proliferative progenitor population characterized by a pronounced chondrogenic bias and limited adipogenic potential.

### U-CPCs exhibit chondrogenic-biased differentiation and BMP2-enhanced cartilage matrix formation

To determine whether U-CPCs exhibit a distinct differentiation profile compared to SSPC populations derived from other tissues, bone marrow-, adipose-, and dental pulp-derived SSPCs (designated B-SSPCs, A-SSPCs, and D-SSPCs, respectively) were isolated by fluorescence-activated cell sorting (FACS) based on canonical human SSPC surface immunophenotypes. Following FACS enrichment (**Figure S5**), U-CPCs, B-SSPCs, A-SSPCs, and D-SSPCs were subjected to lineage-specific induction conditions, along with parallel non-inductive controls, to assess their osteogenic, chondrogenic, and adipogenic differentiation capacities. After 4 days of culture, Alcian blue staining revealed that U-CPCs exhibited the greatest GAG deposition among all groups, even in the absence of chondrogenic induction. D-SSPCs showed moderate GAG deposition, whereas B-SSPCs and A-SSPCs showed minimal staining. Following chondrogenic induction, U-CPCs maintained superior GAG matrix production and exhibited a greater capacity to form cartilage spheroids than other SSPC populations (**Figure 2A**). Osteogenic differentiation was assessed using ARS staining. No detectable mineralized nodules were observed in any of the four populations under noninductive conditions. Following osteogenic induction, B-SSPCs formed the most abundant mineralized nodules, whereas D-SSPCs formed more mineralized nodules than U-CPCs and A-SSPCs, but fewer than B-SSPCs (**Figure 2B**). In contrast, U-CPCs displayed a limited adipogenic potential. Oil Red O staining revealed no detectable lipid accumulation in any population under non-inductive conditions. Following adipogenic induction, B-SSPCs, A-SSPCs, and D-SSPCs exhibited varying degrees of intracellular lipid droplet formation, whereas U-CPCs contained fewer lipid droplets than the other SSPC populations (**Figure 2C**). Together, these findings demonstrate that U-CPCs exhibit a pronounced chondrogenic bias, accompanied by relatively limited osteogenic and adipogenic capacities.

**Figure 2.**
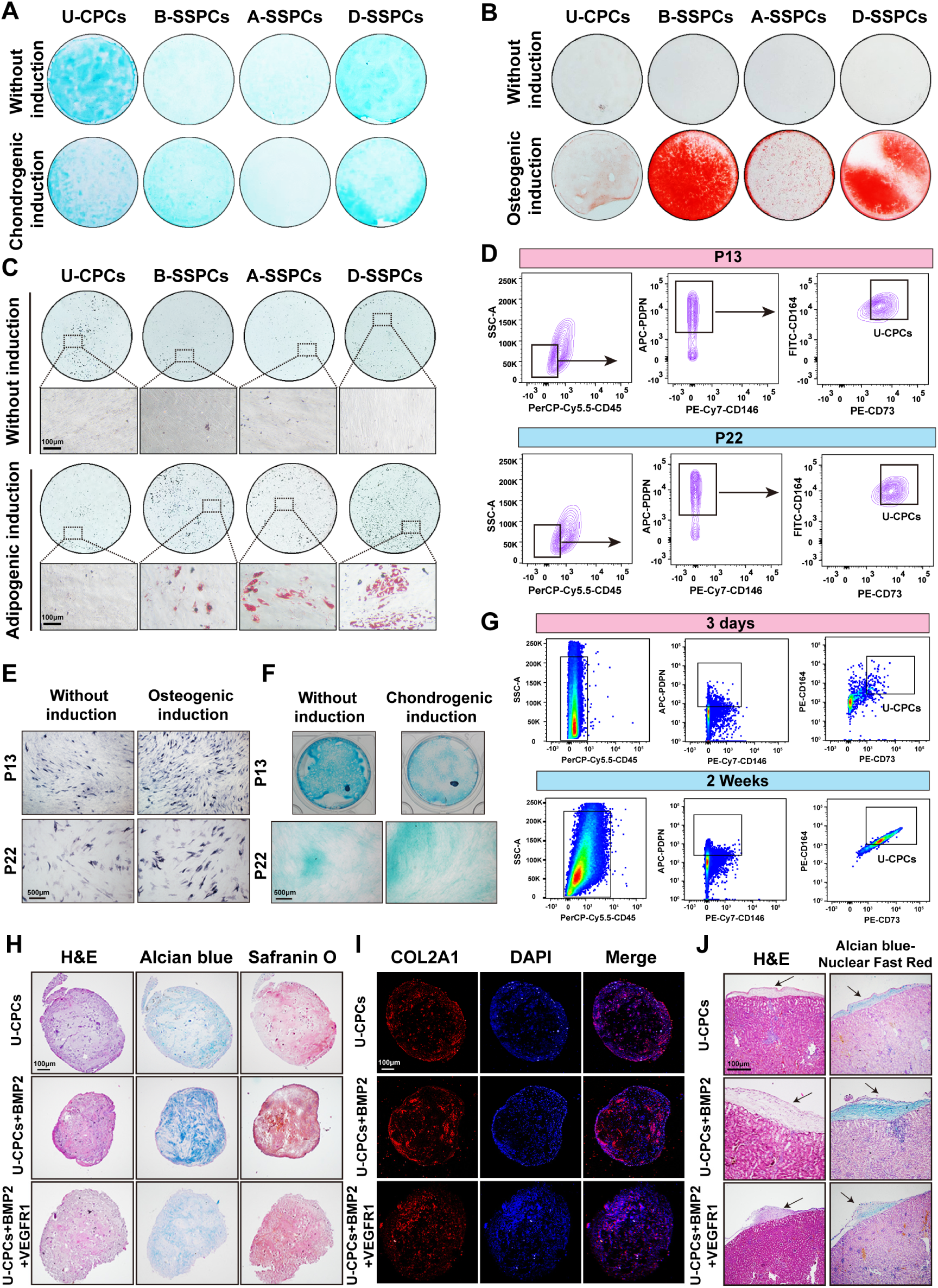
Characterization of U-CPC differentiation potential, phenotypic stability, and cartilage matrix formation *in vitro* and *in vivo*. (A) Alcian blue staining to assess the chondrogenic differentiation of U-CPC or SSPC from different sources with or without chondrogenic induction. (B) ARS staining to evaluate the osteogenic differentiation of the indicated cells under basal or osteogenic-induced conditions. (C) Oil Red O staining to assess the adipogenic differentiation of U-CPC or SSPC from different sources cultured with or without adipogenic induction. (D) Flow cytometric analysis of U-CPCs marker expression during prolonged passaging from P13 to P22. (E and F) Assessment of osteogenic and chondrogenic differentiation of U-CPCs during prolonged passaging by ALP (E) and Alcian blue staining (F). (G) Flow cytometric analysis of the expression of U-CPCs retrieved at 3 days and 2 weeks after subcutaneous implantation. (H) H&E, Alcian blue, and Safranin O staining of Matrigel-embedded U-CPCs treated with BMP2 or BMP2 plus VEGFR1 *in vitro*. (I) Immunofluorescence staining for COL2A1 in Matrigel-embedded U-CPCs treated with BMP2 or BMP2 plus VEGFR1 *in vitro*. (J) H&E and Alcian Blue-Nuclear Fast Red staining of renal capsule implants containing Matrigel-encapsulated U-CPCs treated with BMP2 or BMP2 + VEGFR1.

Next, we assessed whether this cellular phenotype was maintained during prolonged *in vitro* expansion. Flow cytometric analysis of U-CPCs at intermediate (P13) and late (P22) passages showed that their characteristic surface phenotype (CD45⁻ PDPN⁺CD146⁻CD73⁺CD164⁺) was largely preserved at both passages, with no obvious differences between P13 and P22 (**Figure 2D**). However, extended passaging affected their differentiation capacity following lineage-specific induction. Under osteogenic induction, both P13 and P22 cells retained positive ALP activity, although ALP staining was weaker in P22 cells than in P13 cells (**Figure 2E**). Under chondrogenic conditions, P13 cells formed chondrosphere-like structures under both non-inductive and inductive conditions, accompanied by evident GAG deposition. In contrast, P22 cells failed to form obvious chondrosphere-like aggregates, although limited GAG deposition was detectable (**Figure 2F**). These findings indicate that prolonged *in vitro* expansion does not markedly alter the surface phenotype of U-CPCs but progressively compromises their multilineage differentiation capacity.

The phenotypic persistence of U-CPCs was evaluated *in vivo*. Cells were mixed with Matrigel and subcutaneously implanted into immunodeficient nude mice, and the implants were harvested on day 3 and 2 weeks after implantation for flow cytometric analysis. Cells recovered from the implants retained the characteristic U-CPC immunophenotype and were detectable at both time points (**Figure 2G**). The proportion of this population increased from 0.31 ± 0.21% on day 3 to 4.69 ± 1.51% at 2 weeks (**Figure S6**), indicating that U-CPCs can persist *in vivo* while maintaining their phenotypic identity.

Given the intrinsic chondrogenic properties of U-CPCs, we next examined whether their cartilage matrix-forming capacity could be further enhanced by BMP2 and VEGFR1, two factors previously implicated in MSC-mediated cartilage repair ^25^. U-CPCs were evaluated using Matrigel-based *in vitro* culture and *in vivo* renal subcapsular implantation models. After 4 weeks of *in vitro* culture, H&E staining revealed rounded cell-Matrigel aggregates with organized cellular distribution across all groups, whereas Alcian blue and Safranin O staining demonstrated cartilage-like matrix deposition. BMP2 supplementation increased the matrix-positive areas detected by both staining methods compared to U-CPCs alone, whereas additional supplementation with VEGFR1 did not further enhance the BMP2-mediated effect (**Figure 2H**). Quantitative analysis confirmed that the U-CPCs + BMP2 group exhibited the largest Alcian blue- and Safranin O-positive areas among all groups (**Figures S7A and S7B**). Consistently, COL2A1 immunofluorescence intensity was highest in the U-CPCs + BMP2 group (**Figure 2I and Figure S7C**).

Cartilage matrix formation was further examined *in vivo* by implanting U-CPC-containing Matrigel constructs beneath the renal capsules of immunodeficient mice. Four weeks after transplantation, H&E staining revealed matrix-dense ectopic tissue formation within the subcapsular space in all groups, and Alcian Blue-Nuclear Fast Red staining further confirmed the deposition of a GAG-rich matrix. Consistent with the *in vitro* findings, the U-CPCs + BMP2 group exhibited the greatest matrix deposition, followed by the U-CPCs alone group, whereas the U-CPCs + BMP2 + VEGFR1 group showed the least matrix formation among the three groups (**Figure 2J**). This pattern was further supported by a quantitative analysis (**Figure S7D**). These results demonstrate that U-CPCs possess an intrinsic capacity for chondrogenic matrix deposition in both Matrigel-based 3D cultures and *in vivo* implantation models. This matrix-forming capacity was enhanced by BMP2, whereas the addition of VEGFR1 did not further enhance the BMP2-mediated effect.

### U-CPCs show prolonged intra-articular retention and promote cartilage repair in DMM-induced osteoarthritis

To determine whether U-CPCs can persist locally following intra-articular administration, the cells were labeled with the near-infrared fluorescent dye DiR iodide and injected into the knee joints of mice subjected to destabilization of the medial meniscus (DMM). Fluorescence imaging detected DiR signals as early as 6 h post-injection. The signal intensity subsequently increased, peaked at 24 h, and gradually declined thereafter, but remained detectable at all examined time points throughout the 4-week observation period (**Figures 3A and 3B**). To further characterize the tissue-level localization of U-CPCs following intra-articular administration, CM-DiI-labeled cells were injected into the knee joints and examined at 2 and 4 weeks post-injection. CM-DiI-positive signals were detected within the joint tissues at both time points (**Figure S8**), supporting the prolonged local retention of U-CPC-associated fluorescence following intra-articular administration.

**Figure 3.**
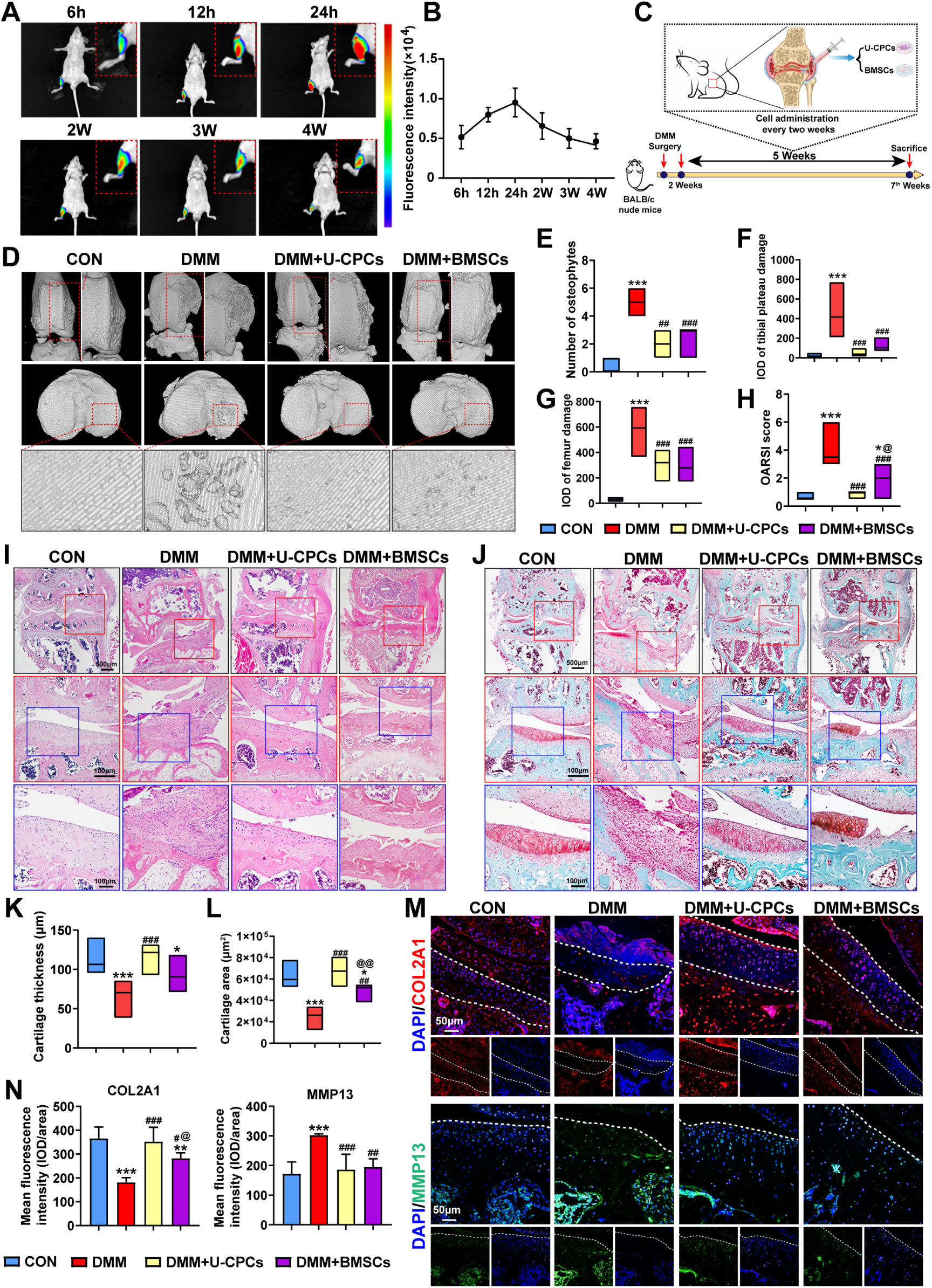
Therapeutic effects of U-CPCs on DMM-induced cartilage injury in mice. (A) Longitudinal *in vivo* fluorescence imaging of DiR-labeled U-CPCs after intra-articular injection. (B) Quantitative analysis of fluorescence intensity after intra-articular injection of DiR-labeled U-CPCs. (C) Schematic illustration of the experimental design and treatment schedule for the DMM-induced cartilage injury model. (D) Representative three-dimensional micro-CT reconstruction images of mouse knee joints after U-CPC or BMSC treatment. (E-G) Quantitative micro-CT analysis of osteophyte number (E) and lesion regions integrated optical density (IOD) at the medial femoral condyle (F), and medial tibial plateau (G). OARSI scores(H) based on Safranin O-Fast Green staining shown in (J). Representative H&E (I) and Safranin O-Fast Green (J) staining of the knee joints after cell treatment. Quantitative analysis of cartilage thickness (K) and cartilage area (L) based on Safranin O-Fast Green staining. (M) Immunofluorescence staining for COL2A1 and MMP13 in the articular cartilage. (N) Quantitative analysis of COL2A1 and MMP13 mean fluorescence intensity (integrated optical density/area). Data are presented as mean ± SD. \**P*<0.05, \*\**P*<0.01, \*\*\**P*<0.001 *vs* CON; ^#^*P*<0.05, ^##^*P*<0.01, ^###^*P*<0.001 *vs* DMM; ^@^*P*<0.05, ^@@^*P*<0.01, ^@@@^*P*<0.001 *vs* DMM+U-CPCs.

Next, we evaluated the therapeutic efficacy of U-CPCs in a DMM-induced cartilage injury model using BALB/c nude mice. Two weeks after DMM surgery, the mice were randomized into four groups: sham-operated control (CON), untreated DMM, DMM + U-CPCs, and DMM + BMSCs. Mice in the two treatment groups received intra-articular injections of U-CPCs or BMSCs, and cartilage repair and joint structural changes were evaluated after five weeks of treatment (**Figure 3C**). Joint swelling was monitored on a weekly basis. One week after treatment, the DMM group exhibited significantly greater joint swelling than the CON group, whereas intra-articular administration of either U-CPCs or BMSCs significantly reduced joint swelling (**Figure S9A**). This effect persisted until week 2 (**Figure S9B**). By week 4, joint swelling subsided across all groups, with no significant differences detected (**Figure S9C**). Thermal nociceptive responses were further assessed using the hot plate test at 2 and 4 weeks after the treatment. U-CPC treatment significantly prolonged withdrawal latency compared with the untreated DMM group at both time points, whereas BMSC treatment showed a similar but weaker trend that did not reach statistical significance (**Figure S10**).

Gross examination revealed severe cartilage surface erosion and prominent osteophyte formation in the medial femoral condyles of untreated DMM joints. These abnormalities were alleviated in both cell-treated groups, which exhibited significantly lower macroscopic OA scores than those of the untreated DMM group (**Figure S11**). Consistently, micro-CT imaging revealed pronounced osteophyte formation and structural damage at the medial femoral condyle and medial tibial plateau in untreated DMM joints, whereas these abnormalities were attenuated by both U-CPCs and BMSCs treatments (**Figure 3D**). Quantitative analysis of reconstructed micro-CT images further showed that both cell treatments significantly reduced osteophyte numbers and lesion-region IOD values at the medial femoral condyle and beneath the medial tibial plateau compared with the untreated DMM group (**Figures 3E-G**).

Histological analysis further supports these structural improvements. H&E staining showed marked osteophyte formation at the medial femoral condyle and medial tibial margin adjacent to the synovial edge in untreated DMM joints, along with severe fibrosis and focal defects of the medial tibial plateau. Intra-articular administration of either U-CPCs or BMSCs suppressed fibrosis, reduced osteophyte formation, and improved cartilage morphology in OA rats. Compared with BMSC-treated joints, U-CPC-treated joints exhibited a more regular cartilage surface and better preservation of chondrocytes at the medial tibial plateau (**Figure 3I**). Safranin O-Fast Green staining revealed weak proteoglycan staining, surface irregularities, and indistinct tidemarks in the DMM group. Both cell treatments restored proteoglycan levels and improved tidemark definition, whereas U-CPC-treated joints displayed a more continuous cartilage matrix than BMSC-treated joints (**Figure 3J**). Quantitative analysis showed that both treatments increased cartilage thickness and area and reduced OARSI scores compared with the untreated DMM group. Notably, U-CPC treatment resulted in significantly greater improvements in the cartilage area and OARSI score than BMSC treatment (**Figures 3H, 3K, and 3L**).

To further assess the quality of cartilage repair, joint sections were immunofluorescently stained with COL2A1 and MMP13 (**Figure 3M**). Compared to CON joints, DMM-induced cartilage injury was associated with a marked reduction in COL2A1 immunoreactivity and a concomitant increase in MMP13 expression. Both U-CPCs and BMSCs treatments increased COL2A1 expression and decreased MMP13 expression. Quantitative analysis further revealed that COL2A1 fluorescence intensity was significantly higher in the U-CPC group than in the BMSC group, whereas MMP13 fluorescence intensity did not differ significantly between the two treatment groups (**Figure 3N**).

Together, these findings demonstrate that U-CPCs exhibit prolonged local retention following intra-articular administration, effectively attenuate DMM-induced cartilage degeneration, improve cartilage structural integrity, and enhance cartilage matrix preservation.

### Therapeutic effects of U-CPCs in a middle–aged rat model of OA

To evaluate the efficacy of U-CPCs under middle–aged pathological conditions, DMM surgery was performed in middle–aged rats as an OA animal model. Intra-articular administration of U-CPCs was performed on postoperative day 3, along with sham-operated and vehicle-treated OA rats as parallel controls. Joint structural changes were evaluated five weeks after treatment **(Figure 4A)**. Micro-CT reconstruction revealed pronounced osteophyte formation and subchondral bone deterioration in the OA group. In contrast, U-CPC treatment preserved joint surface morphology and reduced osteophyte formation **(Figure 4B)**. Quantitative analysis showed that U-CPC treatment significantly alleviated bone erosion of the medial femoral condyle, increased BV/TV, Tb.Th, and reduced Tb.Pf in this region. U-CPC treatment also decreased the osteophyte number compared with that in the untreated OA group **(Figure 4C)**.

**Figure 4.**
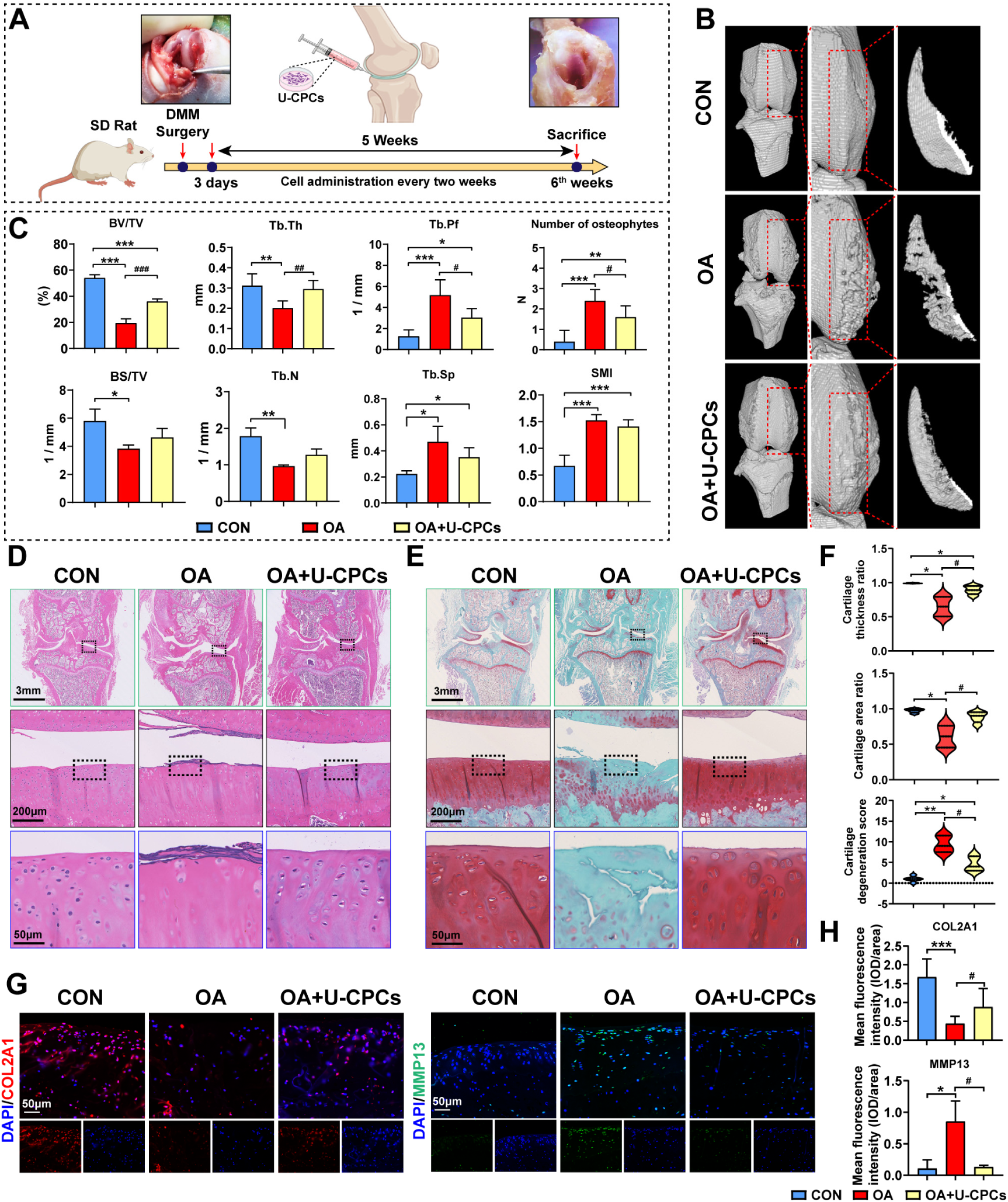
Therapeutic effects of U-CPC treatment in a middle–aged rat model of OA. (A) Schematic of the experimental design. An OA rat model was established in middle–aged rats, followed by intra-articular administration of U-CPCs beginning three days after surgery, repeated every two weeks during the 5-week treatment period. (B) 3D micro-CT of rat knee joints after U-CPC treatment. (C) Quantitative micro-CT analysis of subchondral bone parameters and osteophyte number. (D and E) H&E (D) and Safranin O-Fast Green (E) staining of knee joint sections from each group. (F) Quantitative analysis of cartilage thickness, cartilage area, and cartilage degradation scores based on Safranin O-Fast Green staining. (G) Immunofluorescence staining for COL2A1 and MMP13 in articular cartilage. (H) Quantitative analysis of COL2A1 and MMP13 mean fluorescence intensity. Data are presented as mean ± SD. \**P*<0.05, \*\**P*<0.01, \*\*\**P*<0.001 *vs* CON. ^#^*P*<0.05, ^##^*P*<0.01, ^###^*P*<0.001 *vs* OA.

Histological assessment further supports these findings. H&E staining revealed severe surface fibrosis, chondrocyte loss, and osteophyte formation in OA joints, whereas these degenerative changes were attenuated following U-CPC treatment **(Figure 4D)**. Safranin O-Fast Green staining revealed proteoglycan depletion and poor indistinct tidemarks in the OA group. U-CPC-treated joints exhibited stronger proteoglycan staining and a more clearly defined tidemark structure than the untreated joints **(Figure 4E)**. Quantitative analysis further showed that U-CPCs significantly increased cartilage thickness and area while reducing cartilage degradation scores compared with the OA group **(Figure 4F)**.

To assess extracellular matrix remodeling, immunofluorescence staining was performed for COL2A1 and MMP13 **(Figure 4G)**. OA joints showed significantly diminished COL2A1 fluorescence and increased MMP13 fluorescence. U-CPC treatment effectively reversed this pathological shift, restoring COL2A1 and suppressing MMP13 signals. Quantitative analysis further confirmed these changes **(Figure 4H)**.

Collectively, these findings indicate that U-CPCs attenuate OA-associated structural degeneration and help preserve cartilage matrix homeostasis in middle–aged rats.

### Single-cell RNA-seq and ATAC-seq reveal heightened chondrogenic potential in U-CPCs

To compare the cellular composition and transcriptional states of U-CPCs and UC-MSCs, we performed single-cell RNA sequencing using the 10x Genomics platform (**Figure 5A**). The integrated dataset comprised 19,764 high-quality cells, which were classified into nine MSC-related subpopulations using unsupervised clustering (**Figure 5B**). Although UC-MSCs and U-CPCs were distributed within the same UMAP space, they displayed distinct distributions across clusters and differed markedly in their subpopulation compositions (**Figure 5C and 5D**). The marker gene expression patterns further validated the annotated subpopulations (**Figure 5E**). GO enrichment analysis showed that extracellular matrix organization-related terms were enriched across several MSC subpopulations, whereas cartilage-related terms, including cartilage development and chondrocyte differentiation, were most prominently enriched in the U-CPC-derived EDN1⁺ MSC subpopulation (**Figure 5F**). Accordingly, U-CPCs expressed higher levels of chondrogenic and growth plate-associated genes, including SOX9 and GDF5 (**Figure 5G**). GSVA further demonstrated higher enrichment scores for gene sets related to growth plate cartilage development and chondrocyte differentiation in U-CPCs than in UC-MSCs (**Figure 5H**). Collectively, these findings indicate that U-CPCs exhibit a transcriptional program biased toward chondrogenic differentiation.

**Figure 5.**
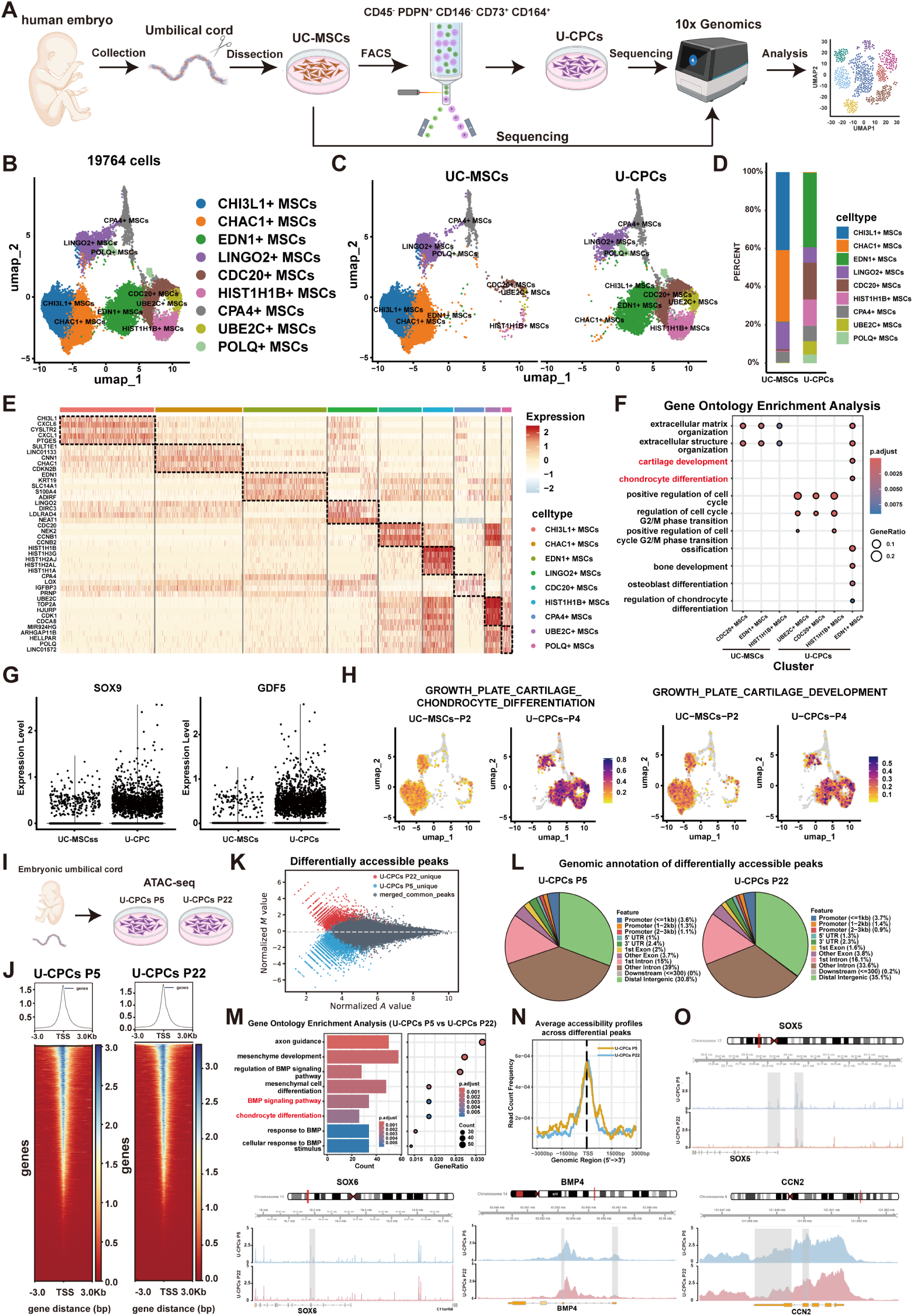
Single-cell RNA-seq and ATAC-seq analysis of U-CPCs. (A) Experimental workflow for the isolation of UC-MSCs from human embryonic umbilical cord tissue, enrichment of U-CPCs by FACS, and single-cell RNA sequencing using the 10x Genomics platform. (B) UMAP visualization of 19,764 cells identified nine MSC-related subpopulations. (C) Distribution of UC-MSCs and U-CPCs in the integrated UMAP space. (D) Relative proportions of the nine MSC-related subpopulations in UC-MSCs and U-CPCs. (E) Heatmap showing the expression of marker genes used to annotate MSC-related subpopulations. (F) GO enrichment analysis of the marker genes from each subpopulation. (G) SOX9 and GDF5 expression in UC-MSCs and U-CPCs. (H) GSVA of gene sets associated with growth plate cartilage development and chondrocyte differentiation in P2 UC-MSCs and P4 U-CPCs. (I) Experimental design of ATAC-seq analysis using early- and late-passage U-CPCs (P5 and P22, respectively). (J) Transcription starts site enrichment profiles used to assess the ATAC-seq library quality. (K) Differential chromatin accessibility analysis between P5 and P22 U-CPCs. (L) Genomic annotation of differentially accessible regions. (M) GO enrichment analysis of genes associated with differentially accessible regions. (N) Aggregate chromatin accessibility profiles across different accessible regions. (O) Genome browser tracks showing chromatin accessibility at the SOX6, BMP4, and CCN2 loci in P5 and P22 U-CPCs. Figure 5A was created using BioRender and has confirmed publishing and licensing rights, with agreement number LS2A5F00BS.

To determine whether this molecular feature is associated with passage-dependent (aging) changes in chromatin accessibility, we performed ATAC-seq on early- and late-passage U-CPCs (P5 and P22, respectively; **Figure 5I**). Both samples showed pronounced transcription start site (TSS) enrichment, supporting the quality of ATAC-seq libraries (**Figure 5J**). Differential accessibility analysis identified several peaks that differed between P5 and P22 cells (**Figure 5K**). These regions were predominantly located in the promoter-proximal, intronic, and distal intergenic regions (**Figure 5L**). GO analysis of genes associated with differentially accessible regions between P5 and P22 identified significant enrichment of biological processes related to chondrocyte differentiation, mesenchymal development, and BMP signaling (**Figure 5M**). Aggregate profiles across the differential peaks further confirmed passage-associated differences in chromatin accessibility (**Figure 5N**). Consistent with this, genome browser tracks showed greater accessibility near chondrogenesis-associated loci, including SOX6, BMP4, and CCN2, in P5 cells than in P22 cells (**Figure 5O**). Together, these findings indicate that serial passaging is associated with epigenetic remodeling of chondrogenic and BMP-related regulatory programs in U-CPCs.

### U-CPC-Derived sEVs Drive Chondrocyte Maturation and Matrix Synthesis

Given the therapeutic effects of U-CPCs in cartilage repair, we examined whether U-CPC-derived small extracellular vesicles (U-CPC-sEVs) contribute to their interaction with chondrocytes. sEVs were isolated from the conditioned medium of U-CPCs using differential ultracentrifugation. TEM revealed that the isolated U-CPC-sEVs exhibited a typical rounded morphology with a clear bilayer membrane structure (**Figure 6A**). NTA showed that most sEVs were distributed within the 30-150 nm range, with a mean diameter of approximately 75 nm (**Figure 6B**). Western blot analysis further demonstrated the enrichment of the canonical sEVs markers CD9 and CD63, together with the absence of the endoplasmic reticulum marker calnexin (**Figure 6C**), confirming the successful isolation of U-CPC-sEVs.

**Figure 6.**
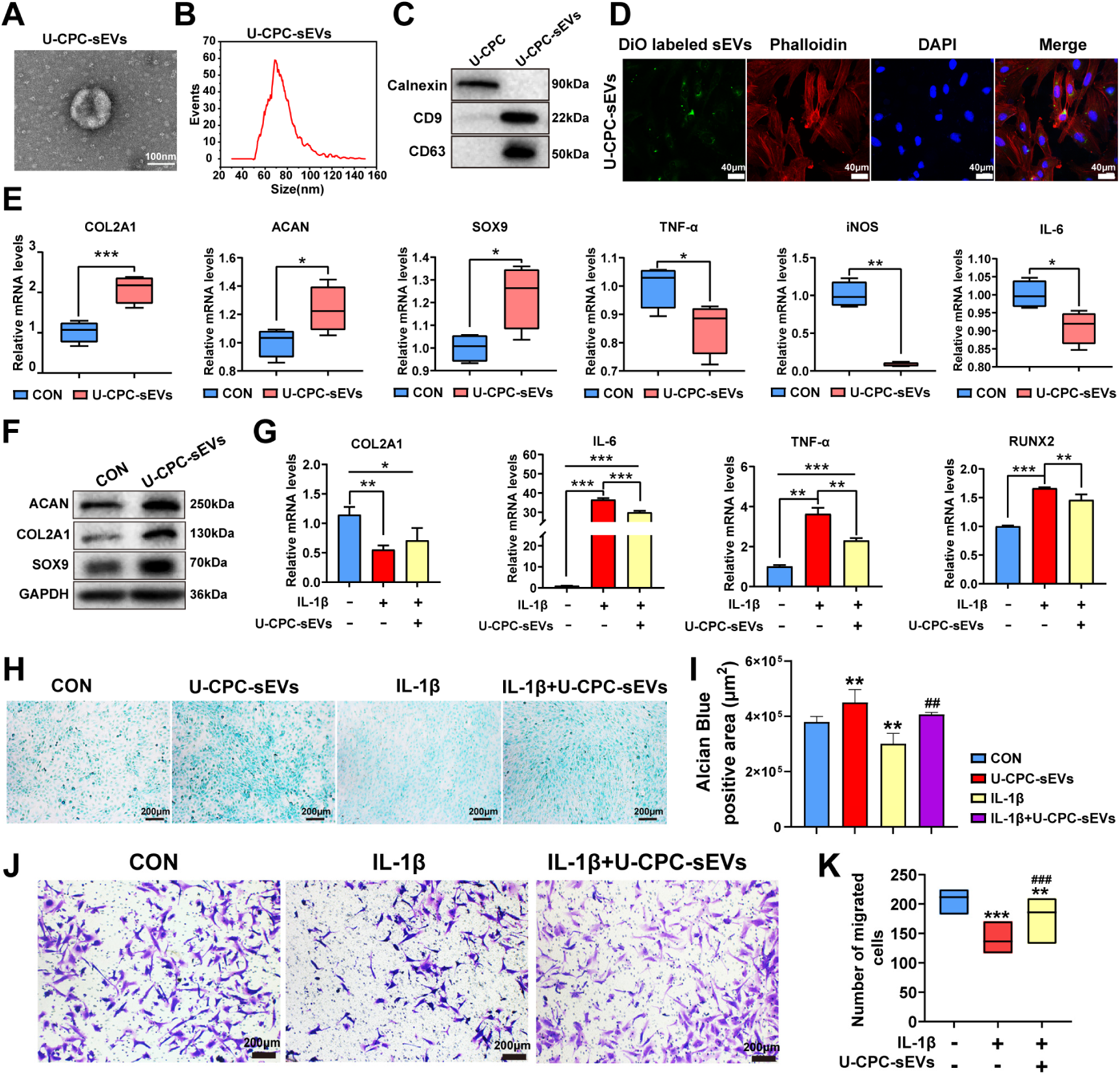
Characterization of U-CPC-sEVs and their effects on chondrocyte activity and cartilage matrix formation. (A) TEM images of U-CPC-sEVs. (B) Particle size distribution of U-CPC-sEVs determined by NTA; (C) Western blot analysis of the sEVs markers (CD63 and CD9) and negative control (Calnexin) in U-CPCs and U-CPC-sEVs. (D) Uptake of DiO-labeled U-CPC-sEVs by C28 Chondrocytes. DiO-labeled sEVs are shown in green, phalloidin-stained cytoplasm in red, and DAPI-stained nuclei in blue. (E) RT-qPCR analysis of chondrogenic genes (*COL2A1, ACAN*, and *SOX9*) and inflammatory markers (*TNF-α, iNOS,* and *IL-6*) following U-CPC-sEV treatment. (F) Western blot analysis of COL2A1, ACAN, and SOX9 expression following U-CPC-sEV treatment. (G) RT-qPCR analysis of *COL2A1, IL-6, TNF-α* and *RUNX2* expression in IL-1β-stimulated C28 chondrocytes after treatment with U-CPC-sEVs. (H) Alcian Blue staining of GAG-rich matrix deposition treated with U-CPC-sEVs under basal or IL-1β-induced inflammatory conditions. (I) Quantitative analysis of GAG-rich matrix deposition based on Alcian Blue staining. (J) Representative images of Transwell migration assays under IL-1β-induced inflammatory conditions with or without U-CPC-sEV treatment. (K) Quantitative analysis of migrated chondrocytes from the Transwell migration assay. Data are presented as mean ± SD. \**P*<0.05, \*\**P*<0.01, \*\*\**P*<0.001 *vs* CON; ^#^*P*<0.05, ^##^*P*<0.01, ^###^*P*<0.001 *vs* IL-1β induced group.

Next, we evaluated whether the U-CPC-sEVs could be internalized by the recipient chondrocytes. Purified U-CPC-sEVs were labeled with the green fluorescent dye, DiO and co-incubated with chondrocytes for 12 h. Fluorescence microscopy revealed a punctate green signal distributed within the phalloidin-stained cytoplasm (**Figure 6D**), indicating the efficient internalization of U-CPC-sEVs by chondrocytes.

Prior to functional evaluation, CCK-8 analysis showed that U-CPC-sEVs enhanced chondrocyte viability under both basal and IL-1β-induced inflammatory conditions, with 6 μg/mL being identified as the optimal concentration for subsequent experiments (**Figure S12**).

Next, we investigated whether U-CPC-sEVs regulate the anabolic phenotype of C28 chondrocytes *in vitro.* Under basal culture conditions, C28 chondrocytes were treated with U-CPC-sEVs for 72 h. RT-qPCR analysis revealed that U-CPC-sEV treatment significantly upregulated the expression of key chondrogenic genes (*COL2A1*, *ACAN*, and *SOX9*) and downregulating pro-inflammatory markers (*TNF-α*, i*NOS*, and *IL-6*) compared to the untreated control group (**Figure 6E**). Consistent with these transcriptional changes, western blot analysis showed increased COL2A1, ACAN, and SOX9 protein levels following U-CPC-sEV treatment (**Figure 6F**), indicating that U-CPC-sEVs promote an anabolic phenotype in chondrocytes.

To further assess whether U-CPC-sEVs could protect chondrocytes under inflammatory conditions, C28 chondrocytes were stimulated with IL-1β for 72 h. IL-1β stimulation markedly suppressed *COL2A1* expression and elevated the expression of hypertrophic (*RUNX2*) and inflammatory (*IL-6* and *TNF-α*) genes compared to the control group. Notably, co-treatment with U-CPC-sEVs significantly rescued *COL2A1* expression and suppressed *RUNX2* and *TNF-α* levels compared to the IL-1β-treated group (**Figure 6G**), suggesting that U-CPC-sEVs alleviate IL-1β-induced inflammatory and phenotype-disruptive responses in chondrocytes.

Next, we assessed GAG-rich matrix deposition using Alcian Blue staining. Under basal conditions, U-CPC-sEV treatment increased GAG-rich matrix deposition compared to that in untreated control cells. In contrast, IL-1β stimulation markedly reduced alcian blue staining, whereas U-CPC-sEV treatment partially rescued GAG-rich matrix deposition under inflammatory conditions (**Figure 6H**). Quantification of the Alcian blue-positive area showed consistent results (**Figure 6I**), supporting that U-CPC-sEVs enhanced the matrix-forming capacity of chondrocytes and mitigated the inhibitory effects of IL-1β on GAG-rich matrix deposition.

Finally, we examined whether U-CPC-sEVs improved chondrocyte migration under inflammatory conditions. Transwell assays showed that IL-1β stimulation markedly reduced the number of migrated chondrocytes, whereas co-treatment with U-CPC-sEVs significantly increased chondrocyte migration compared to IL-1β treatment alone (**Figures 6J and 6K**). These findings suggest that U-CPC-sEVs alleviate IL-1β-induced impairment of chondrocyte matrix synthesis and migration.

### Let-7a-5p promotes chondrogenesis and suppresses hypertrophy by targeting DUSP7 and regulating MAPK signaling

To identify the bioactive miRNA cargo underlying U-CPC-sEV-mediated cartilage repair, we performed small RNA sequencing to profile the miRNAs enriched in U-CPC-sEVs. The five most abundant miRNAs were miR-127-3p, miR-100-5p, let-7a-5p, miR-143-3p, and miR-222-3p (**Figure 7A**). To assess their potential roles in chondrogenic regulation, synthetic mimics of these five miRNAs were transfected into C28 chondrocytes, and successful overexpression was confirmed using RT-qPCR (**Figure 7B**). At 24 h after transfection, all five miRNA mimics significantly increased *SOX9* mRNA expression compared to the negative control (NC) mimic (**Figure 7C**). However, western blot analysis at 96 h showed that only let-7a-5p markedly increased COL2A1 and SOX9 protein expression, whereas the other candidate miRNAs exerted limited or no detectable effects (**Figure 7D**), identifying let-7a-5p as a prominent functional miRNA candidate in U-CPC-sEVs.

**Figure 7.**
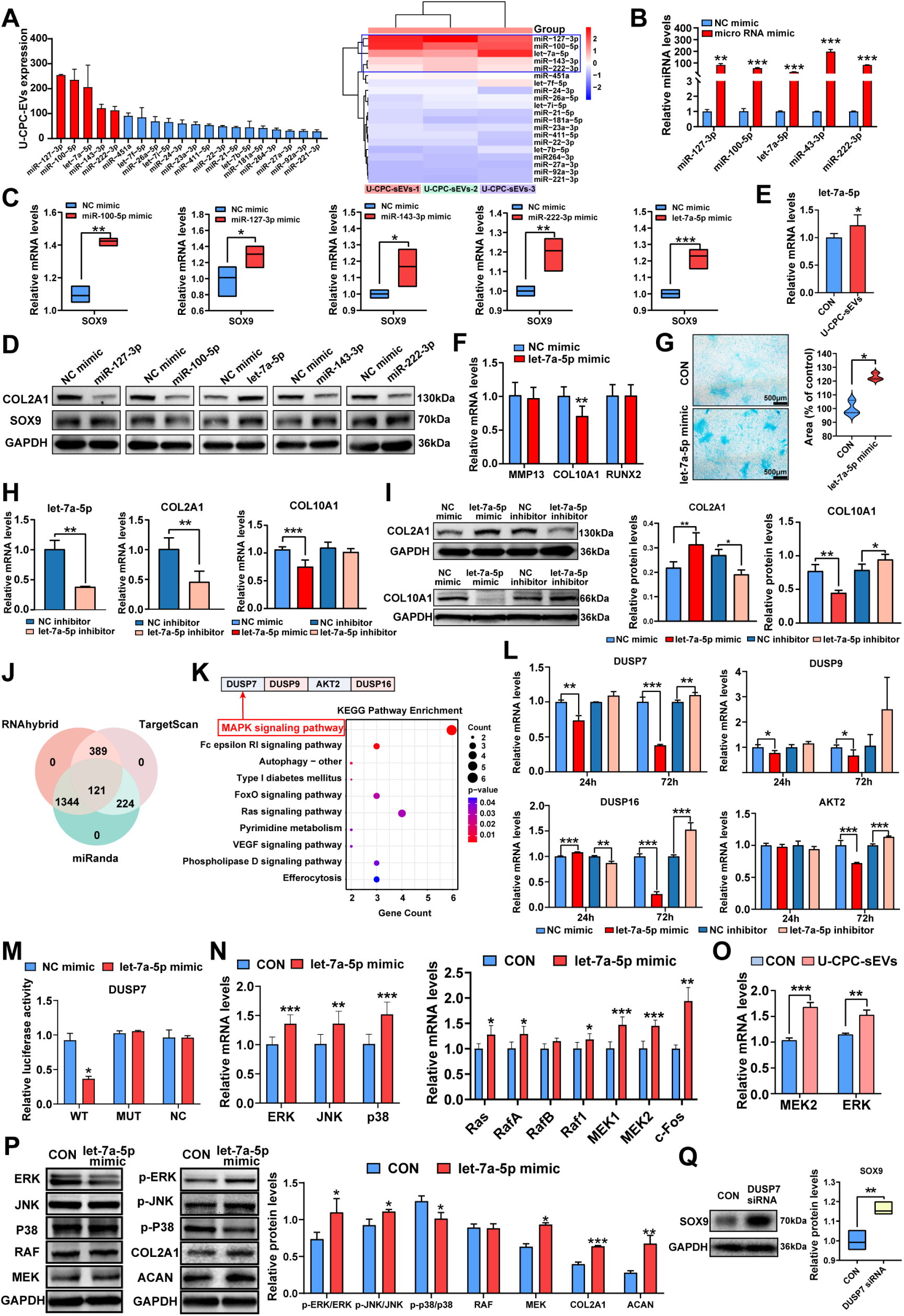
U-CPC-sEVs-delivered let-7a-5p regulates chondrocyte activity by targeting DUSP7 and modulating MAPK signaling. (A) Small RNA sequencing analysis of miRNAs in U-CPC-sEVs. (B) RT-qPCR validation of five candidate miRNA mimics in C28 chondrocytes. (C) RT-qPCR analysis of *SOX9* expression after transfection with the five candidate miRNA mimics. (D) Western blot analysis of COL2A1 and SOX9 expression after treatment with the five candidate miRNA mimics. (E) RT-qPCR analysis of let-7a-5p levels after U-CPC-sEVs treatment. (F) RT-qPCR analysis of *MMP13, COL10A1,* and *RUNX2* expression after let-7a-5p mimic transfection. (G) Alcian Blue staining and quantification of GAG-rich matrix deposition in let-7a-5p mimic-treated cells. (H) RT-qPCR analysis of let-7a-5p, *COL2A1*, and *COL10A1* expression after treatment with a let-7a-5p mimic or inhibitor. (I) Western blot analysis and corresponding quantification of COL2A1 and COL10A1 after let-7a-5p mimic or inhibitor treatment. (J) Prediction of potential let-7a-5p target genes using the RNAhybrid, TargetScan, and miRanda databases. (K) KEGG pathway enrichment analysis of predicted target genes of let–7a–5p. (L) RT-qPCR validation of *DUSP7, DUSP9, DUSP16,* and *AKT2* expression after let-7a-5p mimic or inhibitor treatment. (M) Dual-luciferase reporter assay of let-7a-5p targeting *DUSP7*. (N) RT-qPCR analysis of MAPK pathway-related genes (*ERK, JNK, p38, Ras, RafA, RafB, Raf1, MEK1, MEK2,* and *c-Fos*) after let-7a-5p mimic transfection. (O) RT-qPCR analysis of *MEK2* and *ERK* after U-CPC-sEVs treatment. (P) Western blot analysis of MAPK pathway activity, COL2A1, and ACAN expression after let-7a-5p mimic treatment. (Q) Western blot analysis of SOX9 expression after DUSP7 knockdown. The expected molecular weights of the antibodies are listed as follows: COL2A1(130 kDa), SOX9(70 kDa), COL10A1 (66 kDa), ERK (42-44 kDa), p-ERK (42-44 kDa), JNK (54 kDa), p-JNK (54 kDa), P38 (41 kDa), p-P38 (41 kDa), RAF (75kDa), MEK (43 kDa), ACAN (250 kDa), and GAPDH (36 kDa). Data are presented as mean ± SD (n=3 per group). \**P*<0.05, \*\**P*<0.01, \*\*\**P*<0.001.

Next, we confirmed the delivery of let-7a-5p by U-CPC-sEVs. U-CPC-sEV treatment significantly increased intracellular let-7a-5p abundance in the recipient C28 chondrocytes (**Figure 7E**). To define its effects on the chondrocyte phenotype, we examined markers associated with hypertrophy and matrix degradation. Transfection with the let-7a-5p mimic significantly reduced *COL10A1* mRNA expression without altering *RUNX2* or *MMP13* levels (**Figure 7F**), indicating the selective suppression of the hypertrophic phenotype. Consistently, Alcian blue staining revealed increased GAG-rich matrix deposition in the let-7a-5p group compared with that in the NC group (**Figure 7G**). Complementary loss-of-function experiments supported these findings. The let-7a-5p inhibitor effectively reduced intracellular let-7a-5p levels and decreased *COL2A1* transcript abundance in chondrocytes. Notably, whereas the let-7a-5p mimic reduced *COL10A1* mRNA expression, let-7a-5p inhibition did not significantly alter *COL10A1* transcription (**Figure 7H**). At the protein level, the let-7a-5p mimic increased COL2A1 expression and decreased COL10A1 expression, whereas let-7a-5p inhibition produced the opposite pattern, reducing COL2A1expression and increasing COL10A1 expression (**Figure 7I**). These findings indicate that let-7a-5p promotes an anabolic chondrocyte phenotype while restricting hypertrophic differentiation.

To identify the downstream targets and signaling pathways regulated by let-7a-5p, we integrated predictions from the RNAhybrid, TargetScan, and miRanda databases, yielding 121 overlapping candidate genes (**Figure 7J**). KEGG pathway enrichment analysis identified MAPK signaling as the most enriched pathway. Among the MAPK-associated candidates, *DUSP7*, *DUSP9*, *AKT2*, and *DUSP16* were selected for further validation (**Figure 7K**). RT-qPCR analysis showed that the let-7a-5p mimic significantly reduced *DUSP7* and *DUSP9* mRNA levels at 24 h after transfection, whereas let-7a-5p inhibition produced non-significant upward trends. At 72 h, mimic-mediated suppression of *DUSP7* and *DUSP9* became more pronounced; however, only *DUSP7* expression was significantly increased following let-7a-5p inhibition. *DUSP16* displayed time-dependent opposing responses, whereas *AKT2* was unchanged at 24 h but exhibited a *DUSP7*-like pattern at 72 h post-treatment. Among these candidates, *DUSP7* exhibited the most consistent reciprocal regulation by let-7a-5p (**Figure 7L**). A dual-luciferase reporter assay was performed to further confirm that *DUSP7* is a direct target of let-7a-5p (**Figure 7M**).

We further examined whether let-7a-5p modulates MAPK signaling in C28 chondrocytes. RT-qPCR analysis showed that the let-7a-5p mimic upregulated multiple components of the MAPK cascade, including *Ras*, *RafA*, *Raf1*, *MEK1*, *MEK2*, *c-Fos*, *ERK*, *JNK*, and *p38* (**Figure 7N**). Consistent with these findings, treatment of C28 chondrocytes with U-CPC-sEVs significantly increased MEK2 and ERK expression levels (**Figure 7O**). Immunoblotting further showed that the let-7a-5p mimic increased the phosphorylation ratios of ERK and JNK (p-ERK/ERK and p-JNK/JNK), enhanced MEK activation, and reduced the p-p38/p38). In parallel, let-7a-5p mimic treatment increased the expression of cartilage matrix proteins COL2A1 and ACAN (**Figure 7P**).

Given that let-7a-5p is highly enriched in U-CPC-sEVs and functions as their primary bioactive cargo, these findings provide evidence that U-CPC-sEVs regulate the MAPK cascade in a let-7a-5p-dependent manner. While we established DUSP7 as a direct target of let-7a-5p, miRNAs typically exert pleiotropic effects by silencing multiple transcripts. To determine whether let-7a-5p promotes chondrogenesis primarily through *DUSP7* repression, we performed targeted knockdown experiments. Notably, *DUSP7* knockdown significantly increased SOX9 protein abundance, partially phenocopying the anabolic effects of let-7a-5p (**Figure 7Q**). These results suggest that let-7a-5p promotes an anabolic chondrocyte phenotype, at least in part, by targeting the DUSP7*/* MAPK/ERK signaling axis.

## Discussion

Identifying a reliable cell source for cartilage repair remains a critical clinical hurdle ^26^. Here, we provide compelling evidence that genuine, highly homogenous SSC-like progenitors are not exclusively restricted to skeletal niches. The identification of scalable, lineage-restricted progenitors (U-CPCs) from the umbilical cord expands the current framework of SSC biology. Single-cell profiling demonstrates that U-CPCs are intrinsically lineage-primed-expressing core chondrogenic regulators (SOX9, COL2A1, ACAN) even in the naive state. Consequently, these cells function as committed downstream progenitors, effectively bypassing the functional heterogeneity of conventional MSCs while retaining the expansive capacity necessary for clinical applications. Functionally, these U-CPCs exhibit a robust chondrogenic capacity that surpasses both conventional MSCs and marker-matched SSPCs, sustaining a stable phenotype that resists hypertrophy and lineage drift. Therapeutically, intra-articular administration of U-CPCs attenuates osteoarthritis progression and promotes hyaline cartilage regeneration. Furthermore, we trace this regenerative efficacy beyond cellular engraftment, mapping it directly to the targeted regulation of chondrocytes through a specific sEVs/let-7a-5p/DUSP7/MAPK/ERK signaling cascade.

Tissue-specific epigenetic memory frequently limits the efficacy of standard MSC therapies ^27, 28^. Unlike BMSCs and B-SSPCs, which inherently default to osteogenesis, U-CPCs demonstrate robust chondrogenic plasticity with minimal spontaneous hypertrophy. Mechanistically, this advantage may stem from the avascular, hypoxic environment of the umbilical matrix (Wharton’s jelly), which physiologically preconditions U-CPCs for the harsh articular joint space ^29^. Our transcriptomic data confirm that this hypoxia-adapted phenotype is an intrinsically programmed biological state rather than a transient in vitro artifact, emphasizing the value of isolating lineage-primed subsets from perinatal tissues ^30^. Crucially, our comparative analysis across diverse tissues demonstrates that shared surface immunophenotypes ^11^ do not override the epigenetic memory conferred by the tissues of origin. Despite uniform surface marker expression, B-SSPCs retained a robust osteogenic bias, whereas D-SSPCs displayed a pronounced adipogenic propensity that risks deleterious intra-articular adipose deposition. In contrast, A-SSPCs exhibited restricted capacities for both osteogenesis and chondrogenesis. Collectively, these findings establish that surface markers alone are insufficient predictors of the *in vivo* lineage fate and highlight the profound influence of epigenetic memory on progenitor cell fate.

Consistent with this lineage specification, U-CPCs exhibited greater chondrogenic capacity than bulk BMSCs and UC-MSCs *in vitro*, as reflected by the enhanced deposition of sulfated glycosaminoglycans and type II collagen. Transcriptomic profiling further supported a stable chondroprogenitor state characterized by increased expression of lineage-determining transcriptional programs, including SOX9-associated networks and BMP-related signaling, and reduced osteogenic and adipogenic signatures. Single-cell RNA sequencing additionally revealed substantial intra-population organization within U-CPCs, encompassing a relatively primitive and highly proliferative subset characterized by EDN1, UBE2, CDC20, and HIST1H1B expression, as well as a more differentiated matrix-producing population marked by ACAN and COL2A1expression. By maintaining an equilibrium between proliferative expansion and chondrogenic commitment within this cellular hierarchy, U-CPCs emerge as a distinct and clinically relevant candidate for hyaline cartilage repair.

The regenerative potential of U-CPCs was further confirmed by their therapeutic activity in distinct in osteoarthritis models. Following intra-articular administration, U-CPC-associated signals remained detectable within the joint for more than four weeks, providing a potential basis for sustained local activity. In the DMM mouse model, U-CPC treatment improved nociceptive responses, reduced osteophyte formation, limited articular surface damage, and preserved cartilage structure and matrix composition. Similarly, in a DMM-induced middle-aged OA rat model, U-CPC treatment attenuated cartilage degeneration and preserved subchondral bone microarchitecture. These findings support the therapeutic applicability of U-CPCs in both mechanically induced and degenerative joint disease settings.

Mechanistically, the regenerative activity of U-CPCs appears to be mediated, at least in part, through a potent paracrine program involving small extracellular vesicles (sEVs), consistent with the broader concept that stromal cells can exert therapeutic effects through the secretion of bioactive factors ^31, 32^. U-CPC-derived sEVs were sufficient to reproduce key pro-chondrogenic and protective effects, and small RNA sequencing identified let-7a-5p as a prominent functional miRNA cargo associated with these responses. Although let-7a-5p has been extensively studied as a tumor-suppressive miRNA involved in cellular metabolism ^33, 34^, emerging evidence has also implicated let-7a-5p in tissue regeneration, immunomodulation, and control of cellular senescence during joint repair ^35, 36, 37^. Its functional target network and downstream signaling pathways stay largely uncharacterized. Rather than restricting our investigation to the phenotypic effects of U-CPC-sEVs-delivered microRNA let-7a-5p in OA, we systematically delineated its specific target genes and mechanisms of action.

Consequently, we identified cytoplasmic phosphatase DUSP7, a key mitogen-activated protein kinase phosphatase, as a direct target of let-7a-5p. DUSP7 has been shown to act as a negative regulator of the MAPK signaling pathway ^38, 39^. By suppressing DUSP7, let-7a-5p relieves this inhibitory constraint and promotes MAPK signaling in chondrocytes. It co-activates the Ras-Raf-MEK-ERK and JNK cascades, which drive c-Fos transcription, chondrocyte proliferation, and early matrix maturation ^40, 41^. In parallel, dampening p38 phosphorylation may attenuate chondrocyte catabolism and hypertrophic differentiation, given that activated p38 signaling upregulates matrix-degrading enzymes, including MMP13, and promotes chondrocyte apoptosis ^42^. Moreover, given the established role of p38 activation in promoting osteogenic differentiation ^43^, the reduced p-p38/p38 ratio following let-7a-5p treatment suggests a concomitant restraint on hypertrophy and endochondral ossification in chondrocytes. This coordinated signaling pattern is accompanied by increased COL2A1 and ACAN expression and reduced hypertrophic differentiation, thereby favoring the preservation and synthesis of the cartilage matrix ^44^.

Together, U-CPC-derived sEVs actively modulate recipient chondrocyte behavior through the delivery of let-7a-5p rather than functioning solely as passive vesicular carriers ^45, 46, 47^. By directly targeting DUSP7 and reshaping downstream MAPK signaling, let-7a-5p promotes an anabolic chondrogenic phenotype while restricting hypertrophic and catabolic responses. Thus, the let-7a-5p/DUSP7/MAPK/ERK axis provides a mechanistic link between the lineage-intrinsic properties of U-CPCs and their capacity to promote joint homeostasis and cartilage regeneration.

Limitations of the Study: First, the in vivo efficacy of U-CPCs was evaluated exclusively in rodent models. Given the differences in joint biomechanics, cartilage thickness, and load-bearing profiles between rodents and humans, subsequent validation in large animal models (e.g., porcine or ovine) is required to accurately assess translational potential. Second, although our in vitro data definitively map the U-CPC-sEV-delivered miRNA let-7a-5p to regulate DUSP7/MAPK/ERK axis as a core mechanism for matrix restoration, the detailed spatiotemporal characterization of this signaling cascade within the complex in vivo joint microenvironment warrants further investigation. Translating these mechanistic insights into a standalone, sEV-based cell-free therapeutic modality therefore remains an objective for our subsequent studies.

### Summary

The therapeutic efficacy of bulk UC-MSCs in osteoarthritis remains constrained by cellular heterogeneity, which accounts for variable chondrogenic potential and undesirable hypertrophic drift. To circumvent these limitations, we integrated single-cell transcriptomic profiling with functional screening to resolve UC-MSC heterogeneity, thereby isolating a molecularly defined chondrogenic progenitor subpopulation and delineating the core signaling cascade driving its robust chondrogenesis and articular protection. Unlike conventional unsorted populations, U-CPCs exhibit robust chondrogenic capacity without hypertrophic tendencies while maintaining high proliferative kinetics and an epigenetically youthful state ^48^. Functionally, U-CPCs drive joint homeostasis primarily through targeted sEV secretion. These sEVs endogenously deliver miRNA let-7a-5p, which actively reprograms recipient chondrocyte signaling via the let-7a-5p/DUSP7/MAPK/ERK regulatory axis. Compared with prior studies that mostly applied unseparated UC-MSCs for cartilage repair, our work shifts from heterogeneous bulk cell therapy toward precision progenitor-subset therapy for degenerative joint diseases. These findings not only expand our understanding of MSC subpopulations but also provide a mechanistic basis to develop optimized cell products for future OA regenerative treatment.

## Experimental Sections

### Isolation of CPC by fluorescence-activated cell sorting

Human umbilical cord mesenchymal stem cells (UC-MSCs) and bone marrow mesenchymal stem cells (BMSCs) were isolated according to previously established protocols ^49, 50^, and detailed characterization procedures are provided in the **Supplementary Materials (Figure S1)**. Adipose-derived stem cells (ADSCs) and dental pulp stem cells (DPSCs) were obtained from Cyagen Biosciences (China). All cells were grown in α-Minimum Essential Medium (α-MEM) containing 10% fetal bovine serum (FBS, Gibco, USA) and 1% penicillin-streptomycin (Gibco). To isolate specific skeletal progenitor fractions, the four MSCs were dissociated into single-cell suspensions and subsequently immunolabeled for 30 min at room temperature. As previously reported^11^, a multiplexed fluorophore-conjugated antibody panel was used to target validated human SSC surface markers, including CD45 (PerCP/Cyanine5.5, 1:400; BioLegend, USA), CD146 (PE-Cy7, 1:400, BioLegend), PDPN (APC, 1:400, Thermo Fisher, USA), CD73 (FITC, 1:400, BioLegend), and CD164 (PE, 1:400, BioLegend). After incubation, the strained cells were pelleted by centrifugation, washed, resuspended in fresh α-MEM, and filtered through a 40-μm mesh to ensure a single-cell suspension. Fluorescence-activated cell sorting (FACS) was performed using a BD FACSAria II flow cytometer (BD Biosciences, USA) equipped with an 85-μm nozzle. The resulting SSC-marker-positive subpopulations were designated based on their primary cell source: U-CPCs (derived from UC-MSCs), B-SSPCs (from BMSCs), A-SSPCs (from ADSCs), and D-SSPCs (from DPSCs). Following isolation, the purified progenitor cohorts were cultured and expanded in α-MEM supplemented with 10% FBS and 1% penicillin-streptomycin. All cell lines were maintained at 37℃ in a humidified 5% CO_2_ atmosphere with regular medium changes. Sorting purity was assessed by flow cytometry and analyzed using Flow Jo software (version 10.8.1).

### Proliferation assays

#### CCK-8 assay

UC-MSCs and U-CPCs were seeded in 96-well plates at a density of 1 × 10^3^ cells/well. After 24 h and 48 h of culture, cell proliferation was determined using the CCK-8 assay (Dojindo, Japan), and the absorbance of each well was measured at 450 nm using a multifunctional EPOCH microplate reader (BioTek, USA).

#### Colony formation assay

UC-MSCs and U-CPCs were seeded in six-well plates at a density of 3 × 10^4^ cells/well. After 10 days of culture, the resulting colonies were fixed with methanol, stained with 0.1% crystal violet (Solarbio, China), photographed under a stereomicroscope (RuiHoge, China), and analyzed using Image J (version 1.53k).

### Multilineage differentiation potential of U-CPCs, UC-MSCs, BMSCs, B-SSPCs, A-SSPCs, and D-SSPCs *in vitro*

#### Chondrogenic differentiation

All cell groups were seeded separately in 6-well plates at 1×10^5^ cells/well in complete α-MEM. Upon reaching approximately 90% confluence, the cells were switched to chondrogenic induction medium (OriCell, China), whereas cells maintained in α-MEM without chondrogenic induction served as the non-induced controls. The medium was changed every two days. After 14 days of induction, the cells were washed with PBS, fixed with 4% paraformaldehyde for 30 min, and stained with Alcian Blue (Solarbio) overnight. Stained samples were visualized under an IX73 inverted microscope (Olympus, Japan), and images were analyzed using ImageJ software.

#### Osteogenic Differentiation

All cell groups were seeded in six-well plates at a density of 1×10^5^ cells per well in complete α-MEM. When the cells reached ∼90% confluence, the medium was replaced with osteogenic induction medium (OriCell). The cells were maintained in complete α-MEM without osteogenic induction as non-induced controls. The medium was changed every three days. Alkaline phosphatase (ALP) activity was evaluated on day 7 using a BCIP/NBT ALP Color Development Kit (Beyotime, China). Matrix mineralization was assessed using Alizarin Red S (ARS) staining (OriCell) on day 15. Stained samples were visualized under an inverted microscope, and positively stained areas were quantified using ImageJ software.

#### Adipogenic Differentiation

All cell groups were seeded in six-well plates at a density of 1 × 10^5^ cells per well in complete α-MEM. Upon reaching ∼90% confluence, the medium was replaced with adipogenic induction medium (OriCell). Cells maintained in complete α-MEM without adipogenic induction were used as controls. The medium was changed every three days. After 17 days of induction, lipid accumulation was assessed using Oil Red O staining (Solarbio), according to the manufacturer’s protocol. Stained cells were imaged under an inverted microscope, and the lipid droplet areas were quantified using the ImageJ software.

### Chondrogenic and osteogenic differentiation assessment of long-term expanded U-CPCs

To assess the effect of prolonged in vitro expansion on the differentiation capacity of U-CPCs, cells at passage 13 (P13) and passage 22 (P22) were compared. Osteogenic and chondrogenic potentials were evaluated after 7 and 14 days of differentiation induction via ALP and Alcian blue staining. In parallel, flow cytometry was performed to examine whether the characteristic U-CPCs immunophenotype was maintained across both passages using a defined surface marker panel at a dilution of 1:400.

### Subcutaneous Matrigel implantation for ectopic chondrogenesis and U-CPCs persistence analysis

BMSCs, UC-MSCs, and U-CPCs were individually suspended in ice-cold Matrigel (Corning, USA) at a final density of 5 × 10^3^ cells/μL. For each group, a 250 μL aliquot of the cell-Matrigel suspension was injected subcutaneously into the dorsal neck region of 9∼10 weeks-old male BALB/c nude mice.

For histological analysis, the implants were harvested 4 weeks after transplantation (n = 3 per group), fixed in 4% paraformaldehyde, embedded in paraffin, and sectioned to a thickness of 5 μm. Sections were stained with Hematoxylin and Eosin staining (H&E, Servicebio, China), Alcian Blue, and Safranin O-Fast Green (Solarbio) to evaluate ectopic cartilage-like matrix formation. Histological images were acquired using an ECLIPSE microscope (Nikon, Japan).

For *in vivo* cell tracking, U-CPC-laden Matrigel implants were harvested on day 3 and 2 weeks after transplantation. The retrieved implants were rinsed with PBS, minced into small pieces under sterile conditions, and enzymatically digested with 0.25% trypsin at 37°C to obtain single-cell suspensions. Digestion was terminated by the addition of complete α-MEM, and the suspensions were filtered through a 35-μm cell strainer, centrifuged, and washed. The resulting single-cell suspensions were stained with fluorochrome-conjugated antibodies against U-CPC surface markers, including CD45-PerCP-Cy5.5, CD146-PE-Cy7, PDPN-APC, CD73-FITC, and CD164-PE. Flow cytometry was performed to quantify the proportion of U-CPCs persisting within the retrieved Matrigel implants, and the data were analyzed using FlowJo software.

### Morphogen-directed U-CPCs chondrogenesis in Matrigel-based *in vitro* and renal subcapsular models

To assess whether the selected morphogenic factors regulate U-CPC-mediated cartilage matrix formation, U-CPCs were suspended in ice-cold Matrigel at a density of 3 × 10^4^ cells/μL. For both *in vitro* and *in vivo* assays, Matrigel mixtures were prepared in three groups: (1) U-CPCs in Matrigel alone (U-CPCs group); (2) U-CPCs with BMP2 (50 ng/mL; R&D Systems, USA) within the Matrigel (U-CPCs + BMP2 group); and (3) U-CPCs with BMP2 (50 ng/mL) and vascular endothelial growth factor receptor 1 (VEGFR1; 400 ng/mL, Abcam, UK) within the Matrigel (U-CPCs + BMP2 + VEGFR1 group).

For the *in vitro* 3D culture assay, the prepared Matrigel mixtures were allowed to gel at 37°C and then cultured in α-MEM supplemented with 10% FBS and 1% penicillin-streptomycin in a humidified 5% CO_2_ atmosphere. The cultures were maintained for 4 weeks, and the medium was refreshed every 2 days. The resulting constructs were harvested, fixed, paraffin-embedded and sectioned. The sections were stained with H&E, Alcian blue, and Safranin O-Fast Green to assess tissue architecture and cartilage-like matrix deposition. Immunofluorescence staining for COL2A1 (Novus Biologicals, USA) was performed to evaluate cartilage matrix formation.

For the *in vivo* assay, a renal subcapsular implantation model was used. For each group, 20 μL of the corresponding Matrigel mixture was implanted beneath the left renal capsule of 9-week-old male BALB/c nude mice (n = 3 per group). At 4 weeks post-transplantation, the kidneys containing the implants were harvested, fixed, embedded and sectioned. Ectopic tissue formation and cartilage-like matrix deposition were assessed using H&E and Alcian Blue-Nuclear Fast Red staining.

### *In vivo* tracking of U-CPCs distribution and retention after intra-articular injection

To evaluate the intra-articular retention of U-CPCs, 7-week-old male BALB/c nude mice (n=3) were subjected to destabilization of the medial meniscus (DMM) surgery to induce an osteoarthritic microenvironment, as previously described^51^. Prior to transplantation, U-CPCs were fluorescently labeled with the near-infrared lipophilic dye DiR lodide (Coolaber, China). Following DMM,1×10^6^ labeled cells suspended in 10μL of sterile saline were slowly injected intra-articularly into the operated knees. To assess the biodistribution and retention of U-CPCs, the fluorescent signals of the injected cells were longitudinally monitored at 6, 12, and 24 h, 2 weeks, and 4 weeks post-injection using a multifunctional GelView6000Plus in vivo fluorescence imaging system (BioLight, China).

For tissue-level localization, a separate cohort of DMM-operated mice received intra-articular injections of U-CPCs labeled with the red fluorescent dye CM-DiI (Molecular Probes, USA). Knee joints were harvested at 2 and 4 weeks after the intra-articular injection (n = 3 per time point). The specimens were fixed in 4% paraformaldehyde for 72 h and decalcified in 12% EDTA solution (pH 7.4; Solarbio) for 6 weeks. After decalcification, the joints were embedded in paraffin and sectioned to a thickness of 5 μm. The sections were examined using fluorescence microscopy to determine the localization of CM-DiI-labeled U-CPCs within the joint tissues.

### *In vivo* therapeutic evaluation of U-CPCs in DMM-induced osteoarthritis models *DMM-induced osteoarthritis models and intra-articular treatment*

To evaluate the therapeutic efficacy of U-CPCs in cartilage repair, DMM-induced OA models were established in both 7-week-old male BALB/c nude mice and 32-week-old middle–aged male Sprague-Dawley (SD) rats. For the mouse model, animals were randomly assigned to four groups: sham-operated controls (CON), DMM, DMM + BMSCs, and DMM + U-CPCs (n = 8 per group). Intra-articular treatment was initiated two weeks after DMM surgery. Mice in the treatment groups received 1×10^6^ BMSCs or U-CPCs suspended in 10 µL of 4% albumin (Solarbio), whereas the DMM group received an equal volume of 4% albumin vehicle.

For the middle–aged rat model, animals were randomly assigned to three groups: sham-operated controls (CON), OA, and OA + U-CPCs (n = 7 per group). Intra-articular treatment was initiated three days after the surgery. Rats in the U-CPCs treatment group received 1×10^6^ cells suspended in 100 µL of 4% albumin, whereas rats in the OA group received an equal volume of 4% albumin. All intra-articular injections were administered every two weeks. The animals were euthanized 5 weeks after the first injection, and the knee joints were collected for subsequent histological and molecular analyses.

#### Thermal withdrawal latency assessment

Thermal withdrawal latency was assessed in DMM mice and the corresponding treatment groups at 2 and 4 weeks after intra-articular administration. Nociceptive responses were measured using a hot/cold plate analgesia meter (Panlab, Spain) maintained at 52.5 °C. The latencies to paw withdrawal, licking, and jumping were recorded. A cut-off time of 30 s was applied to prevent thermal injury. Each mouse was tested three times, and the mean withdrawal latency was used for statistical analysis.

#### Micro-CT assessment

The knee joints were scanned using a μCT80 micro-CT system (Scanco Medical, Switzerland). After scanning, the bone images were reconstructed and reoriented in three dimensions using DataViewer software. Quantitative analysis was performed using CTAn software (Version 1.13, Bruker Micro-CT, Belgium) to analyze the erosion on the bone surface of the joint and subchondral bone changes in the knee joint, including bone volume fraction (BV/TV), trabecular thickness (Tb.Th), trabecular number (Tb.N), tissue volume (TV), trabecular pattern factor (Tb.Pf), bone surface-to-tissue volume ratio (BS/TV), trabecular number (Tb.N), trabecular separation (Tb.Sp), and structural model index (SMI). Three-dimensional reconstruction was performed using CTvox software (Bruker, Belgium). Regions of bone damage at the medial femoral condyle and beneath the medial tibial plateau were delineated on reconstructed micro-CT images, and their integrated optical density (IOD; lesion area × mean optical density) was quantified using the Image-Pro Plus software under identical analysis settings.

#### Histological and immunofluorescence assessment

After micro-CT scanning, the knee joint specimens were fixed in neutral-buffered formalin for 72 h and decalcified in 12% EDTA solution for 4 weeks. The samples were then embedded in paraffin and sectioned at 5 µm. Cartilage morphology and matrix preservation were evaluated using H&E and Safranin O-Fast Green staining. The cartilage thickness and area at the medial tibial plateau were quantified using Image-Pro Plus 6.0. Histopathological severity was assessed using the OARSI scoring system for mouse joints and the Mankin-based Histological-Histochemical Grading System (HHGS) for rat joints. For immunofluorescence analysis, the sections were incubated with antibodies against COL2A1 and MMP13 (Signalway Antibody, USA, 1:100), and the images were quantified using ImageJ software. The images were captured under identical exposure settings. The mean fluorescence intensity was quantified using ImageJ software and calculated as the integrated optical density divided by the analyzed area (IOD/area).

### Single-cell RNA and bulk ATAC sequencing analyses

Single-cell RNA sequencing was performed on UC-MSCs and U-CPCs using the 10x Genomics Chromium platform, and bulk ATAC-seq was performed on U-CPCs at P5 and P22. Sequencing data were processed using standard pipelines for quality control, clustering, differential expression, chromatin accessibility, and functional enrichment analysis. Detailed experimental procedures and bioinformatic analyses are described in the **Supplementary Materials**.

### Isolation and characterization of U-CPC-derived sEVs

U-CPCs were maintained in α-MEM consisting of 1% penicillin-streptomycin and 10% sEVs-free FBS (prepared by ultracentrifugation of standard FBS at 170,000 × g for 10 hours at 4 °C, followed by a 0.22-µm filtration). After 48 h of culture, the conditioned medium was harvested and subjected to differential centrifugation at 4 °C to remove dead cells and large debris (300 × g for 10 min; 2,000 × g for 10 min; 10,000 × g for 30 min). The supernatant was filtered (0.22-μm pore size) and subjected to ultracentrifugation at 110,000 × g for 70 min at 4 °C to pellet the sEVs. The pellet was washed with ice-cold PBS, centrifuged again under the same conditions, and resuspended in PBS. The protein concentration of the purified vesicles, namely U-CPC-derived sEVs (U-CPC-sEVs), was determined using a BCA kit (Beyotime, China), following the manufacturer’s instructions. The U-CPC-sEVs were cryopreserved at -80 °C.

For characterization, the morphology and size distribution of sEVs were evaluated using transmission electron microscopy (TEM) and nanoparticle tracking analysis (NTA), respectively. Furthermore, western blotting was performed to confirm the presence of established sEVs markers (CD63 and CD9) and the absence of the intracellular contaminant calnexin.

### Cellular uptake of U-CPC-sEVs

To visualize cellular uptake, U-CPC-sEVs were labeled with 3 μM DiO (Beyotime, China) for 1 h at room temperature. Unbound dye was removed by ultracentrifugation at 110,000 × *g* for 70 min at 4 °C, with the sEVs pellets resuspended in 25 mL PBS between centrifugation steps. Human C28/I2 chondrocytes were incubated with DiO-labeled U-CPC-sEVs (6 μg/mL) for 12 h. After incubation, the cells were washed with PBS to remove non-internalized sEVs, fixed with 4% paraformaldehyde for 40 min, and permeabilized with 0.1% Triton X-100 for 5 min. The F-actin cytoskeleton was stained with Phalloidin-iFluor™ 594 (Abcam, UK), and nuclei were counterstained with DAPI (Invitrogen, USA). Cellular internalization was imaged using an Olympus FV3000 confocal laser scanning microscope (Olympus, Japan), and the fluorescence signals were quantified using ImageJ software.

### RNA isolation and RT-qPCR

Total RNA was extracted from the harvested cells using TRIzol reagent according to a standard phenol-chloroform protocol, as previously described^52^. Complementary DNA (cDNA) was synthesized using the PrimeScript™ RT Reagent Kit (Takara Bio, Japan). Quantitative real-time PCR (qPCR) amplification was performed using an ABI Prism 7500 Real-Time PCR System (Applied Biosystems, USA) with TB Green^®^ Premix Ex Taq™ II (Takara Bio, Japan). Target gene expression was normalized to that of GAPDH.

For miRNA analysis, miR-127-3p, miR-100-5p, let-7a-5p, miR-143-3p, and miR-222-3p, cDNA was synthesized using the RevertAid First Strand cDNA Synthesis Kit (Thermo Scientific, USA) and miRNA-specific stem-loop primers. RT-qPCR was subsequently performed using TB Green^®^ Premix Ex Tap^TM^ Ⅱ, miRNA-specific forward primers, and a universal reverse primer (Sangon Biotech, China). The miRNA expression was normalized to U6 small nuclear RNA. Relative expression levels were calculated using the 2^−ΔΔCT^ method. The primer sequences are listed in **Table S1** in the **Supplementary Materials**.

### Western blot analysis

Total protein from the harvested cells was lysed using RIPA lysis buffer (Thermo Fisher Scientific). Equal amounts of protein (30 μg per sample) were separated by SDS-PAGE and transferred onto PVDF membranes (Merck Millipore, Ireland). After washing with TBST (Solarbio), the membranes were blocked with 5% non-fat milk (Beyotime) and then incubated overnight at 4℃ with primary antibodies against CD63 (Abcam, USA) CD9 (Abcam), Calnexin (Abcam), COL2A1 (Signalway antibody, USA), ACAN (Abcam), SOX9 (Signalway antibody), COL10A1 (Proteintech, China), ERK (Signalway antibody), p-ERK(Abcam), JNK (Signalway antibody), p-JNK (Abcam), p-P38 (Abcam), RAF (Signalway antibody), MEK (Signalway antibody), and GAPDH (Abways, China). All primary antibodies were used at a dilution of 1:1,000. After washing, the membranes were incubated with horseradish peroxidase-conjugated secondary antibodies (1:5000, Proteintech) at room temperature for 1 h, and immunoreactive protein bands were visualized using ECL reagent (Beyotime) and captured using GelView 6000M (Biolight, China).

### MicroRNA (miRNA) sequencing analysis of U-CPC-sEVs

Small RNA sequencing of U-CPC-sEVs was performed by the Beijing Genomics Institute (BGI, China), and downstream analyses were conducted using the BGI Dr Tom system. Potential target genes of the selected miRNAs were predicted using the RNAhybrid, miRanda, and TargetScan databases, followed by KEGG pathway enrichment analysis. Detailed procedures for RNA extraction, library preparation, sequencing, and data processing are provided in **Supplementary Materials**.

### miRNA mimic, inhibitor, and siRNA transfection

Human C28/I2 chondrocytes were seeded into six-well plates at a density of 2× 10^5^ cells/well and cultured for 24 h before transfection. Cells were transiently transfected with the indicated miRNA mimics, including miR-127-3p, miR-100-5p, miR-143-3p, miR-222-3p, let-7a-5p mimic, let-7a-5p inhibitor, or DUSP7 siRNA. The corresponding non-targeting mimics, inhibitors, and siRNA sequences were used as negative controls. Transfection was performed using Lipofectamine 3000 reagent (Invitrogen) according to the manufacturer’s instructions, with a final oligonucleotide concentration of 90 nM. Five hours after transfection, the transfection medium was replaced with complete DMEM supplemented with 10% fetal bovine serum and 1% penicillin/streptomycin. The cells were then cultured for the indicated periods (24 or 72 h) before subsequent molecular analysis. All synthetic RNA oligonucleotides were purchased from Sangon Biotech, and their sequences are listed in **Table S2** in the **Supplementary Materials**.

### Dual-luciferase reporter assay

The wild-type (WT) 3′ untranslated region (3′ UTR) of DUSP7 containing the predicted let-7a-5p binding site and the corresponding mutant (MUT) sequence carrying nucleotide substitutions within the seed-matching region were chemically synthesized and cloned into the pmirGLO Dual-Lucifecrasc miRNA Target Expression Vector by GenePharma (China). For the reporter assay, C28/I2 chondrocytes were seeded in 12-well plates at a density of 5 × 10^5^ cells/well and cultured for 24 h. Cells were then co-transfected with the WT or MUT reporter plasmid together with either the let-7a-5p mimic or a negative control (NC) mimic using Lipofectamine 3000, according to the manufacturer’s instructions. Five hours after transfection, the medium was replaced with fresh complete culture medium. After 48 h, cells were harvested and lysed, and Renilla and Firefly luciferase activities were measured using the Dual-Luciferase Reporter Assay System (Promega, USA) according to the manufacturer’s protocol. Relative luciferase activity was calculated by normalizing the Renilla luciferase activity to the firefly luciferase activity.

## Ethics approval and consent to participate

Animal experiments were authorized by the Academic Committee on Experimental Animal Ethics of Zhanjiang Central Hospital (approval numbers: DW-2025011 and KYSB-2023049). Bone marrow samples and umbilical cord tissues were obtained from donors at the Affiliated Hospital of Guangdong Medical University, with approval from the Ethics Committee of the Affiliated Hospital of Guangdong Medical University (Approval Number: PJGXB2024-001).

## Acknowledgments

This work received financial support from the following sources: the National Nature Science Foundation of China (82604701, 81703584), the Natural Science Foundation of Guangdong Province (2022A1515220166, 2025A1515010196), the Administration of Traditional Chinese Medicine of Guangdong Province, China (20250047), the Science and Technology Foundation of Zhanjiang (2022A01099), and the Discipline Construction Fund of Central People’s Hospital of Zhanjiang (2022A09).

The figure abstract in this article was created using BioRender.com. Publishing license rights were obtained under agreement number EC2A66UH87.

## Authorship contributions

**Xiang Gao:** Investigation, Data management, Software, Data visualization, Formal analysis, Writing-original draft. **Jie Han:** Writing-original draft, Investigation, Data management, Data visualization. **Yankui Zhai:** Investigation, Data Curation, Methodology, Formal analysis. **Jintong Song:** Investigation, Methodology, Formal analysis. **Jia Tian:** Investigation, Methodology, Data management, Software. **Lixian Li:** Investigation, Visualization, Formal analysis. **Dahong Liang:** Investigation, Software, Formal analysis. **Zhiping Chen:** Investigation, Methodology, Visualization. **Jiamei Xu:** Investigation, Methodology. **Yuting Liao:** Investigation, Methodology. **Ahmad**, Validation, Review and editing of the manuscript. **Rui Lin:** Data visualization, Formal analysis. **Liao Cui:** Methodology, Resources. **Bilian Xu:** Resources, Formal analysis. **Li Li:** Resources, Formal analysis. **Qixian Xiao:** Resources, Formal analysis. **Ren Xu:** Supervision, Project administration, Review and editing of the manuscript. **Yanzhi Liu:** Conceptualization, Supervision, Project administration, Funding acquisition, Review and editing of the manuscript.

## Declaration of Generative AI and AI-Assisted Technologies in the Writing Process

During the preparation of this work, the authors used Google Gemini/Deepseek solely to improve language clarity, grammar, and overall readability of the manuscript. After using this tool, the authors critically reviewed and edited the content as necessary and took full responsibility for the final content of the publication. The AI tool was not involved in data generation, analysis, visualization, or the formulation of scientific conclusions. After using these tools, the authors reviewed and edited the content as needed and took full responsibility for the content of the published article.

### Resource availability

#### Lead Contact

Further information and requests for resources and reagents should be directed to and will be fulfilled by the lead contact, Yanzhi Liu

#### Materials Availability

This study did not generate new unique reagents.

#### Data and Code Availability

Single-cell RNA-seq, ATAC-seq, and miRNA-seq data generated in this study have been deposited in the Gene Expression Omnibus (GEO) database under accession number GSE346565. This paper does not report original code. The analyses were performed using publicly available software and referenced packages as described in the STAR Methods. Any additional information required to reanalyze the data reported in this paper is available from the lead contact upon request.

## References

1. Pueyo Moliner, A., et al. Restoring articular cartilage: insights from structure, composition and development. Nat Rev Rheumatol. 21, 291–308 (2025).

2. Randsborg, P. H. et al. Microfracture Versus Arthroscopic Debridement for the Treatment of Symptomatic Cartilage Lesions of the Knee: 2-Year Results From a Multicenter Double-Blinded Randomized Controlled Trial. Am J Sport Med. 53, 2107–2117 (2025).

3. Deng, L. et al. Efficacy and safety of single versus repeated injections of mesenchymal stem cells for the treatment of knee osteoarthritis: a systematic review and network meta-analysis of randomized controlled trials. Stem Cell Res Ther. 16, 505 (2025).

4. Snow, M. et al. A Randomized Trial of Autologous Chondrocyte Implantation Versus Alternative Forms of Surgical Cartilage Management in Patients With a Failed Primary Treatment for Chondral or Osteochondral Defects in the Knee. Am J Sport Med. 51, 367–378 (2023).

5. Li, J. et al. Fibrocartilage hyalinization: A potential therapeutic strategy for articular fibrocartilage. J Orthop Translat. 52, 313–324 (2025).

6. Shi, F. et al. Identification and function of periosteal skeletal stem cells in skeletal development, homeostasis, and disease. J Orthop Translat. 51, 177–186 (2025).

7. Melis, S. et al. Skeletal stem and progenitor cells in bone physiology, ageing and disease. Nat Rev Endocrinol. 21, 135–153 (2025).

8. Yuan, G. et al. Skeletal stem cells in bone development, homeostasis, and disease. Protein Cell. 15, 559–574 (2024).

9. Trompet, D. et al. Skeletal stem and progenitor cells in bone development and repair. Journal of bone and mineral research : the official journal of the American Society for Bone and Mineral Research. 39, 633–654 (2024).

10. Chan, C.K.F. et al. Identification and specification of the mouse skeletal stem cell. Cell. 160, 285–298 (2015).

11. Chan, C.K.F. et al. Identification of the Human Skeletal Stem Cell. Cell. 175, 43–56. e21 (2018).

12. Li, Q. et al. Insights into skeletal stem cells. Bone Res. 10, 61 (2022).

13. Yuan, G. et al. New perspective of skeletal stem cells. Biomater transl. 3, 280–294 (2022).

14. Sun, J, et al. A vertebral skeletal stem cell lineage driving metastasis. Nature. 621, 602–609 (2023).

15. Zhu, Q. et al. Procr^+^ chondroprogenitors sense mechanical stimuli to govern articular cartilage maintenance and regeneration. Cell. 188, 5194–5211.e5116 (2025).

16. Mizuhashi, K. et al. Resting zone of the growth plate houses a unique class of skeletal stem cells. Nature. 563, 254–258 (2018).

17. Zhang, X. et al. Msx1^+^ stem cells recruited by bioactive tissue engineering graft for bone regeneration. Nat Commun. 13, 5211 (2022).

18. Ortinau, L. C. et al. Identification of Functionally Distinct Mx1^+^αSMA^+^ Periosteal Skeletal Stem Cells. Cell Stem Cell. 25, 784–796.e785 (2019).

19. Han, Y. et al. Lkb1 deletion in periosteal mesenchymal progenitors induces osteogenic tumors through mTORC1 activation. J Clin Invest. 129, 1895–1909 (2019).

20. Yang, G. et al. Identification of the metaphyseal skeletal stem cell building trabecular bone. Sci Adv. 10, eadl2238 (2024).

21. Pineault, K. M. et al. Hox11 expressing regional skeletal stem cells are progenitors for osteoblasts, chondrocytes and adipocytes throughout life. Nat Commun. 10, 3168 (2019).

22. Ambrosi, T. H. et al. Human skeletal development and regeneration are shaped by functional diversity of stem cells across skeletal sites. Cell Stem Cell. 32, 811–823.e811 (2025).

23. Josephson, A. M. et al. Age-related inflammation triggers skeletal stem/progenitor cell dysfunction. P Natl Acad Sci USA. 116, 6995–7004 (2019).

24. Ambrosi, T. H. et al. Aged skeletal stem cells generate an inflammatory degenerative niche. Nature. 597, 256–262 (2021).

25. Murphy, M. P. et al. Articular cartilage regeneration by activated skeletal stem cells. Nat med. 26, 1583–1592 (2020).

26. Han, X. et al. Mesenchymal stem cells in treating human diseases: molecular mechanisms and clinical studies. Signal transduct target ther. 10, 262 (2025).

27. Reinisch, A. et al. Epigenetic and in vivo comparison of diverse MSC sources reveals an endochondral signature for human hematopoietic niche formation. Blood. 125, 249–260 (2015).

28. Kfoury, Y. et al. Mesenchymal cell contributions to the stem cell niche. Cell Stem Cell. 16, 239–253 (2015).

29. Ong, L.J.Y. et al. Localized Oxygen Control in a Microfluidic Osteochondral Interface Model Recapitulates Bone–Cartilage Crosstalk During Osteoarthritis. Adv Funct Mater. 34, 2315608 (2024).

30. Zhang, S. et al. Single-cell transcriptome analysis of uncultured human umbilical cord mesenchymal stem cells. Stem Cell Res Ther. 12, 25 (2021).

31. Kou, M. et al. Mesenchymal stem cell-derived extracellular vesicles for immunomodulation and regeneration: a next generation therapeutic tool? Cell Death Dis. 13, 580 (2022).

32. Kumar, MA. et al. Extracellular vesicles as tools and targets in therapy for diseases. Signal transduct target ther. 9, 27 (2024).

33. Shi, Y. et al. Let-7a-5p inhibits triple-negative breast tumor growth and metastasis through GLUT12-mediated warburg effect. Cancer Lett. 495, 53–65 (2020).

34. Duan, S. et al. Exogenous Let-7a-5p Induces A549 Lung Cancer Cell Death Through BCL2L1-Mediated PI3Kγ Signaling Pathway. Front Oncol. 9, 808 (2019).

35. Chen, S. Y. et al. Engineered extracellular vesicles carrying let-7a-5p for alleviating inflammation in acute lung injury. Biomed Sci. 31, 30 (2024).

36. Shao, L. et al. Let-7a-5p derived from parathyroid hormone (1-34)-preconditioned BMSCs exosomes delays the progression of osteoarthritis by promoting chondrocyte proliferation and migration. Stem Cell Res Ther. 16, 299 (2025).

37. Li, Q. et al. Platelet-rich plasma-derived microRNA let-7a-5p alleviates knee osteoarthritis by regulating macrophage polarization and improving inflammatory microenvironment. Front Immunol. 17, 1756467 (2026).

38. Bai, H. et al. DUSP7 inhibits cervical cancer progression by inactivating the RAS pathway. J Cell Mol Med. 25, 9306–9318 (2021).

39. Liu, J. et al. Dual Specific Phosphatase 7 Exacerbates Dilated Cardiomyopathy, Heart Failure, and Cardiac Death by Inactivating the ERK1/2 Signaling Pathway. Cardiovasc Transl Res. 15, 1219–1238 (2022).

40. Ma, N. et al. The regulatory mechanism of p38/MAPK in the chondrogenic differentiation from bone marrow mesenchymal stem cells. J Orthop Surg Res. 14, 434 (2019).

41. Li, J. et al. MEK/ERK and p38 MAPK regulate chondrogenesis of rat bone marrow mesenchymal stem cells through delicate interaction with TGF-β1/Smads pathway. Cell Prolif. 43, 333–343 (2010).

42. Gao, Y. et al. Obacunone acts as a histone deacetylase 1 inhibitor to limit p38MAPK signaling and alleviate osteoarthritis progression. J Orthop Surg Res. 20, 441 (2025).

43. Liu, S. et al. The hDPSC-CM-derived peptide P1DC promotes osteogenic differentiation by regulating the p38/MAPK pathway. Stem Cell Res Ther. (2026).

44. Zhang, W. et al. Upregulated desmin/integrin β1/MAPK axis promotes elastic cartilage regeneration with increased ECM mechanical strength. Int J Biol Sci. 19, 2740–2755 (2023).

45. Wang, J. et al. A synergistic strategy involving reverse-adaptation and engineered MSC exosomes against ferroptosis in osteoarthritis. Biomaterials. 330, 124024 (2026).

46. Piñeiro-Ramil, M. et al. Mesenchymal stromal cells-derived extracellular vesicles in cartilage regeneration: potential and limitations. Stem Cell Res Ther. 16, 11 (2025).

47. Wang, L. et al. Engineering Extracellular Vesicles as Delivery Systems in Therapeutic Applications. Adv Sci (Weinh). 10, e2300552 (2023).

48. Fang, L. et al. Immunophenotypic characterization and clinical relevance of passage number in umbilical cord mesenchymal stromal cells: A review. Cell Transplant. 35, 9636897261433504 (2026).

49. Lee, AY. et al. Minimal Cube Explant Provides Optimal Isolation Condition of Mesenchymal Stem Cells from Umbilical Cord. Tissue Eng Regen Med. 19, 793–807 (2022).

50. Baghaei, K. et al. Isolation, differentiation, and characterization of mesenchymal stem cells from human bone marrow. Gastroenterol Hepatol Bed Bench. 10, 208–213 (2017).

51. Liu, Y. et al. Sargassum polysaccharide attenuates osteoarthritis in rats and is associated with the up-regulation of the ITGβ1-PI3K-AKT signaling pathway. J Orthop Translat. 47, 176–190 (2024).

52. Dymond, JS. Explanatory chapter: quantitative PCR. Methods Enzymol. 529, 279–289 (2013).

