## Supplementary Materials for "A distinct chondrogenic progenitor population from human umbilical cord– derived mesenchymal stromal cells for osteoarthritis therapy"

**Experimental Sections**

**Characterization of UC-MSCs and BMSCs**

Human umbilical cord-derived mesenchymal stem cells (UC-MSCs) and bone marrow-derived mesenchymal stem cells (BMSCs) were harvested by trypsin digestion, followed by centrifugation, and resuspended at a density of 1 × 10^6^ cells/mL. For phenotypic identification, UC-MSCs were incubated with antibodies against CD34, CD45, CD11b, CD19, and human leukocyte antigen-antigen D related (HLA-DR), CD73, and CD90, whereas BMSCs were incubated with antibodies specific for CD45, CD73, CD90, and CD105. Surface marker expression was subsequently analyzed using a FACSCanto II flow cytometer (BD Biosciences, USA) following the manufacturer’s protocol. All antibodies were purchased from BD Biosciences, and the flow cytometry data were processed using FlowJo software (version v10.8.1).

The multilineage differentiation potential of UC-MSCs and BMSCs was evaluated by culturing the cells in osteogenic, adipogenic, and chondrogenic induction media. Osteogenic differentiation was maintained continuously for 14 days, whereas adipogenic and chondrogenic inductions were extended for 21 days. Differentiation was confirmed by Alizarin Red S, Oil Red O, and Alcian Blue staining, respectively.

**Single-cell RNA and bulk ATAC sequencing analyses**

***Single-cell RNA sequencing***

Single-cell RNA sequencing (scRNA-seq) was performed by Guangzhou Kidio Biotechnology Co., Ltd. (China). Single-cell suspensions were loaded onto a 10x Genomics Chromium Controller to generate gel bead-in-emulsions (GEMs). cDNA libraries were prepared using the Chromium Next GEM Single Cell 3' Reagent Kit according to the manufacturer's protocol, and sequenced on an Illumina platform. Raw base call (BCL) files were processed using the Cell Ranger pipeline (version 7.0.1; 10x Genomics) to output gene-barcode expression matrices. Downstream analyses were executed in R utilizing the Seurat package. Cells were filtered using standard quality criteria (nFeature_RNA > 500 and percent.mt < 20), and doublets were removed using DoubletFinder (v2.0). After normalization, scaling, and dataset integration, principal component analysis (PCA) and uniform manifold approximation and projection (UMAP) were used for dimensionality reduction, followed by graph-based clustering and annotation based on marker genes and cluster-specific differentially expressed genes. Differential expression analysis was performed using the Wilcoxon rank-sum test. Gene Ontology (GO) enrichment analysis was conducted using clusterProfiler (v4.18.4), and gene set variation analysis (GSVA; v2.4.4) was performed to evaluate cartilage-related gene sets.

***Bulk ATAC sequencing***

Bulk assay for transposase-accessible chromatin with high-throughput sequencing (ATAC-seq) was performed by Shanghai Jiayin Biotechnology Co., Ltd. (China). Briefly, U-CPCs at P5 and P22 were harvested to map chromatin accessibility dynamics during replicative senescence. Following cell lysis and isolation of intact nuclei, a tagmentation reaction was executed utilizing a hyperactive Tn5 transposase to simultaneously cleave and tag open chromatin regions with sequencing adapters. The resulting DNA fragments were purified, amplified via PCR, and the constructed libraries were sequenced on an Illumina platform. Raw reads were quality-filtered using fastp (v1.0.1), aligned to the human reference genome (GRCh38) using Bowtie2 (v2.5.4). Low-quality reads and PCR duplicates were removed using SAMtools (v1.22.1) and Picard (v3.4.0). Signal tracks and heatmaps were generated using alignmentSieve, bamCoverage, and plotHeatmap. Peaks were called using MACS2 (v2.2.9.1), and differential accessibility between P5 and P22 cells was assessed using MAnorm (v1.1.4). Peaks were annotated using ChIPseeker (v1.46.1) with TxDb.Hsapiens.UCSC.hg38. knownGene, and genes associated with differential peaks were subjected to GO enrichment analysis using clusterProfiler (v4.18.4). Genome browser tracks were generated using Gviz (v1.54.0), with gene annotations retrieved via biomaRt (v2.66.0). Signal tracks and heatmaps were generated using deepTools.

**microRNA sequencing analysis of U-CPC-sEVs**

Total RNA was isolated from U-CPC-sEVs. Its integrity and concentration were assessed using the DNF-471 standard sensitivity RNA analysis kit. Three independent biological replicates were evaluated. Small RNA libraries incorporating Unique Molecular Identifiers (UMIs) were constructed according to the manufacturer’s instructions. and sequenced on the BGI platform (BGI, Shenzhen, China). PCR products ranging from 110 to 130 bp were size-selected by polyacrylamide gel electrophoresis before library quality assessment and sequencing. Raw reads were processed to remove adaptor sequences, low-quality reads, and reads shorter than 15 nucleotides. Clean reads were aligned to the human reference genome using Bowtie2 (v2.2.5), and known and novel miRNAs were annotated using miRDeep2. miRNA abundance was quantified using UMI counts and normalized as follows:

$$\text{Expression}=\frac{C\times{10}^{6}}{T}$$

where $C$represents the UMI count assigned to a specific miRNA and $T$ represents the total number of clean UMI reads in the corresponding sample.

**CCK-8 assay**

C28/I2 chondrocytes were seeded in 96-well plates at a density of 1×10^4^ cells per well and cultured for 24 h. Under basal conditions, cells were treated with U-CPC-sEVs at concentrations of 3,6, and 12 μg/mL for 24 h. For the inflammatory model, cells were exposed to 1 ng/mL IL-1β alone or in combination with the indicated concentrations of U-CPC-sEVs for 24 h. Untreated cells served as the control.

After treatment, the CCK-8 assay was added to each well according to the manufacturer’s instructions, followed by incubation for 2 h at 37 °C. Absorbance was measured at 450 nm using an Epoch microplate reader (BioTek, USA).

**Transwell migration assay**

The migratory capacity of C28/I2 chondrocytesb following EV treatment was evaluated using 24-well Transwell inserts with an 8 μm pore size (Corning, USA). C28/I2 cells were suspended in a serum-free medium and seeded into the upper chambers at 1×10^4^ cells/well. The cells were assigned to three groups: untreated control, 1 ng/mL IL-1β alone, and 1 ng/mL IL-1β combined with 6 μg/mL U-CPC-sEVs. The lower chambers were filled with 600 μL of complete culture medium containing 10% FBS. After 12h of incubation, the Transwell chambers were fixed with 4% paraformaldehyde for 15 min and stained with 0.1% crystal violet (Beyotime, China) for 20 min at room temperature. After thorough PBS rinsing, non-migrated cells remaining on the upper membrane were gently wiped off with cotton swabs. The cells migrated from upper side to the lower side of the chamber were photographed under a DMI 3000B inverted microscope (Leica, Germany). Migrated cells were counted in six randomly selected fields per well using Image J software.

**Results:**

**Characterization of BMSCs and UC-MSCs**

Multilineage differentiation of the isolated BMSCs and UC-MSCs was confirmed histologically: osteogenic induction yielded robust calcium nodule formation, as evidenced by Alizarin Red S staining; chondrogenic induction produced abundant sulfated glycosaminoglycans marked by Alcian Blue, and adipogenic conversion resulted in extensive intracellular lipid accumulation stained by Oil Red O (**Figures S1 A, C**). Flow cytometric analysis further confirmed the characteristic MSCs surface marker profiles of the isolated cells. BMSCs exhibited high expression of CD90 (99.9%), CD105 (99.5%), and CD73 (100%), with minimal expression of CD45 (0.12%). Similarly, UC-MSCs showed high expression of CD90 (100%), CD105 (96.3%), and CD73 (100%), whereas the combined expression of CD45, CD34, CD11b, CD19, and HLA-DR was low (0.91%) (**Figures S1 B, D**). Collectively, these findings confirmed that both BMSCs and UC-MSCs possessed multilineage differentiation capacity and characteristic MSCs immunophenotypes with high purity.

**Supplementary tables and figures**

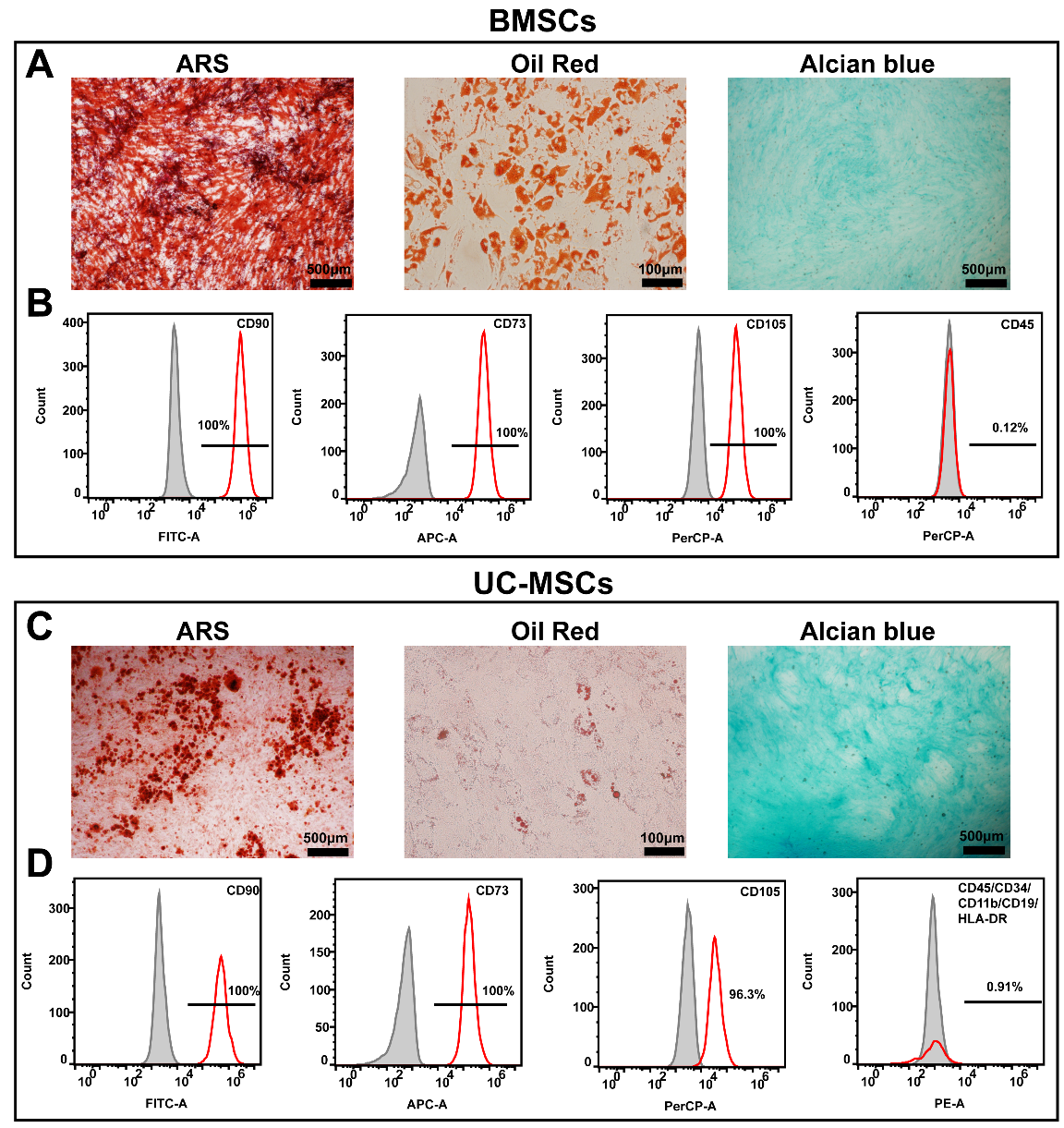

**Figure S1.** Identification of BMSCs and UC-MSCs. (A) Multilineage differentiation of BMSCs toward osteogenic, adipogenic, and chondrogenic lineages, as demonstrated by Alizarin Red (ARS), Oil Red O, and Alcian blue staining, respectively. (B) Flow cytometric analysis showed that BMSCs were positive for CD90, CD73, and CD105 and negative for CD45. (C) Multilineage differentiation of UC-MSCs toward osteogenic, adipogenic, and chondrogenic lineages, as demonstrated by ARS, Oil Red O, and Alcian blue staining, respectively. (D) Flow cytometric analysis showed that UC-MSCs were positive for CD73, CD90, and CD105 and negative for CD19, CD11b, CD34, CD45, and HLA-DR.

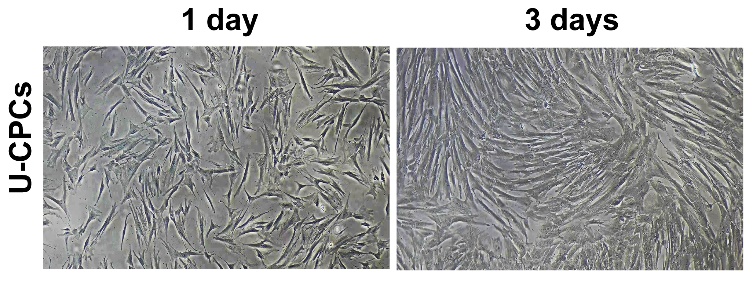

**Figure S2.** Morphology of U-CPCs after 1 day and 3 days of culture.

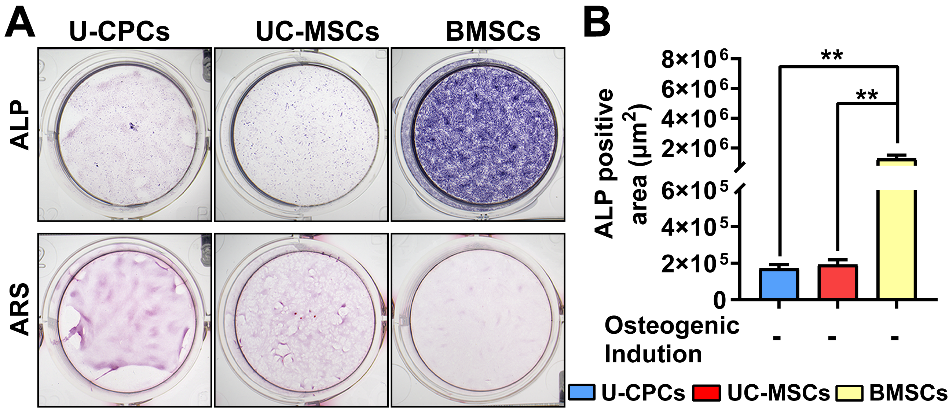

**Figure S3.** Assessment of osteogenic differentiation under non-induced conditions. (A) Representative ALP and Alizarin Red S (ARS) staining of U-CPCs, UC-MSCs, and BMSCs cultured without osteogenic induction. (B) Quantitative analysis of ALP staining under non-osteogenic induction conditions. (C) Representative Oil Red O staining of U-CPCs, UC-MSCs, and BMSCs cultured without adipogenic induction.

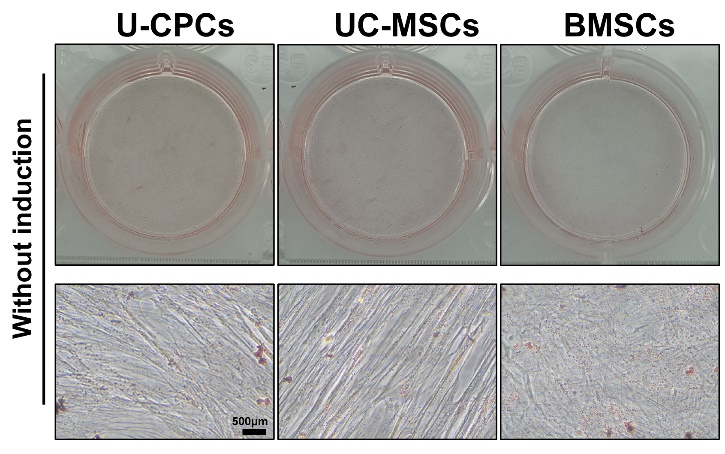

**Figure S4.** Representative Oil Red O staining of U-CPCs, UC-MSCs, and BMSCs cultured without adipogenic induction.

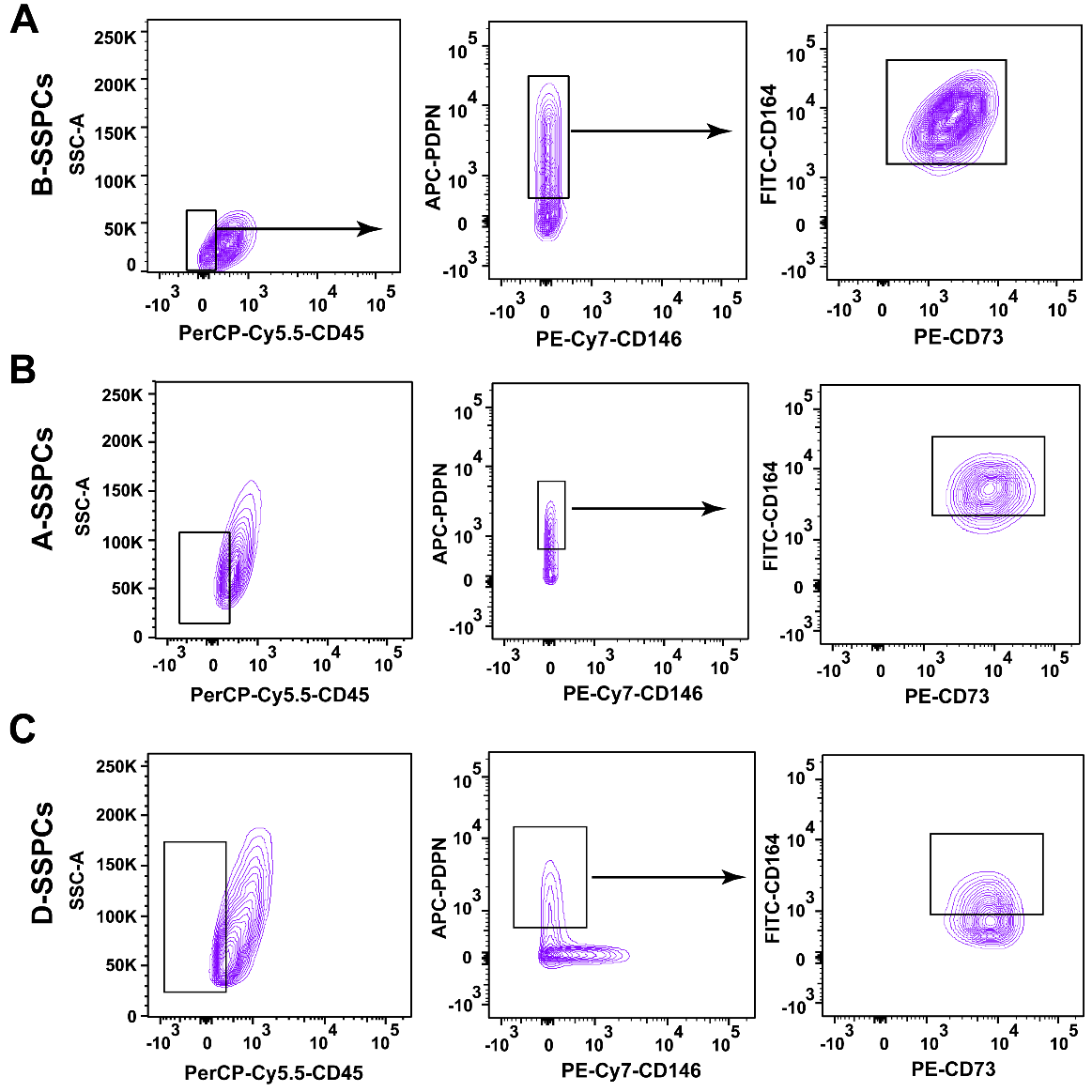

**Figure S5.** FACS isolation of SSPCs. Representative FACS plots showing the isolation of B-SSPCs from BMSCs (A), A-SSPCs from ADSCs (B), and D-SSPCs from DPSCs (C).

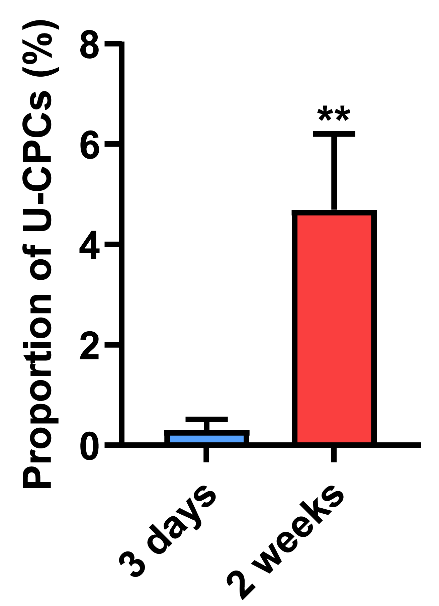

**Figure S6.** Proportion of U-CPCs recovered from subcutaneous implants at day 3 and 2 weeks, as determined by flow cytometry. Data are presented as the mean ± SD. ^**^*P* < 0.01.

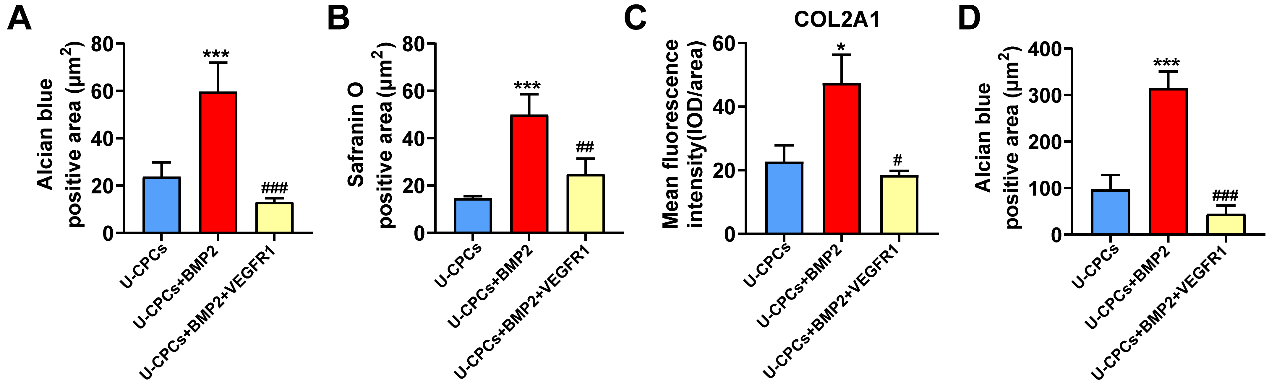

**Figure S7.** Quantitative analysis of chondrogenic differentiation of U-CPCs. Quantification of Alcian blue (A) and Safranin O-positive areas (B) in Matrigel-embedded U-CPCs treated with BMP2 or BMP2 plus VEGFR1 *in vitro*. (C) Quantification of COL2A1 immunofluorescence intensity in Matrigel-embedded U-CPCs treated with BMP2 or BMP2 plus VEGFR1 *in vitro*. (D) Quantification of the Alcian blue-positive area in renal capsule implants containing Matrigel-encapsulated U-CPCs treated with BMP2 or BMP2 plus VEGFR1. Data are presented as mean ± SD. ^*^*P*<0.05, ^**^*P*<0.01, ^***^*P*<0.001 *vs* U-CPCs; ^#^*P*<0.05, ^##^*P*<0.01, ^###^*P*<0.001 *vs* U-CPCs+BMP2.

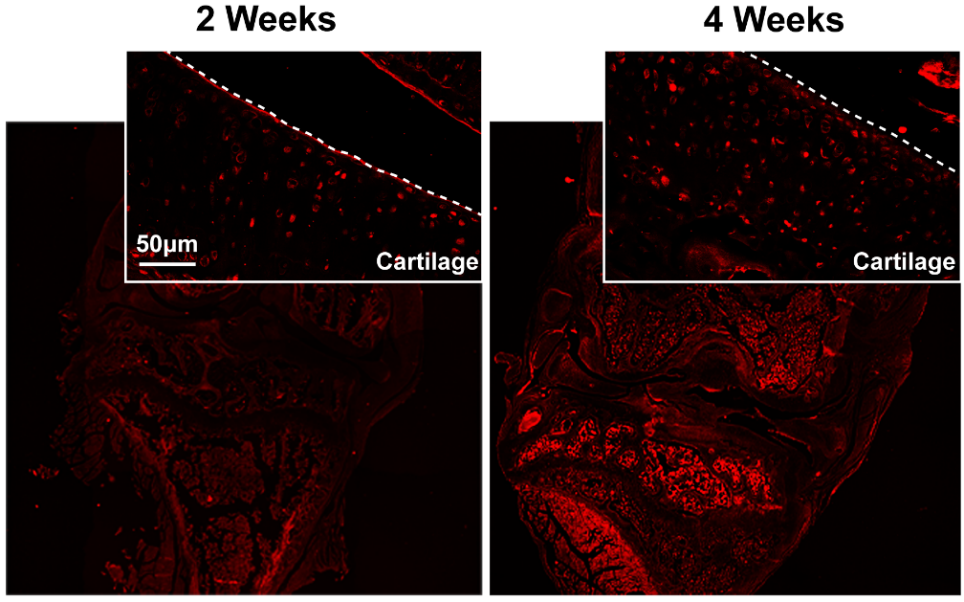

**Figure S8.** Representative confocal images of CM-DiI-labeled U-CPCs in knee joint sections at 2 and 4 weeks post-injection.

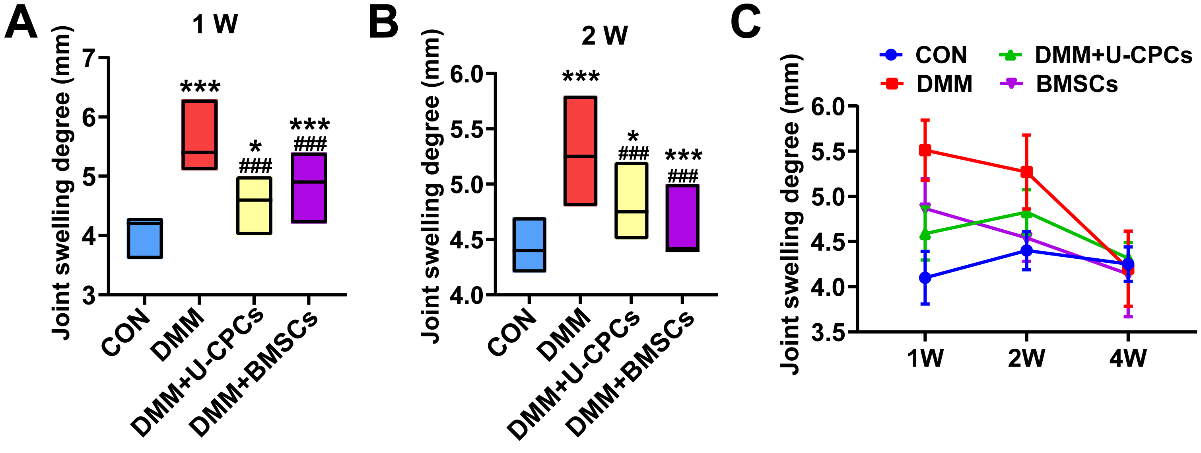

**Figure S9.** Quantification of joint swelling in DMM-induced OA mice receiving intra-articular U-CPCs or BMSCs injection at 1 week (A), 2 weeks (B), and 4 weeks (C). Data are presented as mean ± SD. ^*^*P*<0.05, ^**^*P*<0.01, ^***^*P*<0.001 *vs* CON; ^#^*P*<0.05, ^##^*P*<0.01, ^###^*P*<0.001 *vs* DMM; ^@^*P*<0.05.

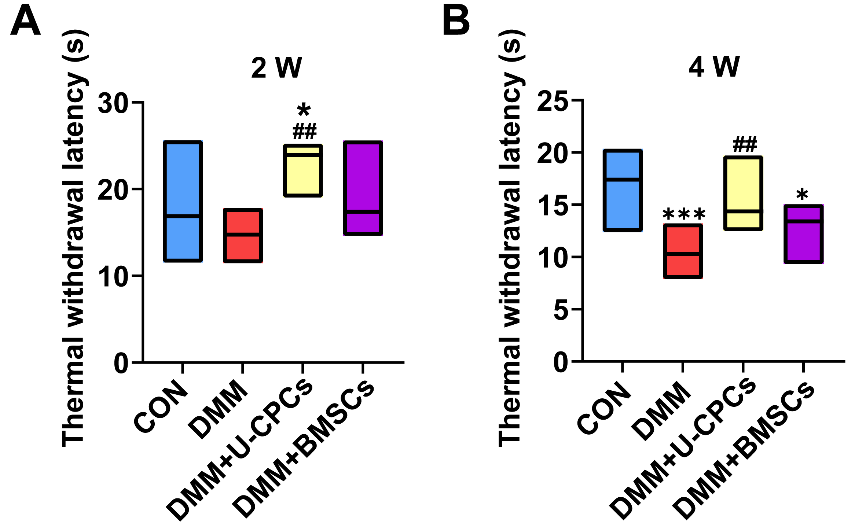

**Figure S10.** Thermal withdrawal latency in mice with DMM-induced osteoarthritis. Withdrawal latency was measured using the hot plate test at 2 weeks (A) and 4 weeks (B) after treatment. Data are presented as mean ± SEM; n = 6 per group. ^*^*P*<0.05, ^**^*P*<0.01, ^***^*P*<0.001 *vs* CON; ^#^*P*<0.05, ^##^*P*<0.01, ^###^*P*<0.001 *vs* DMM.

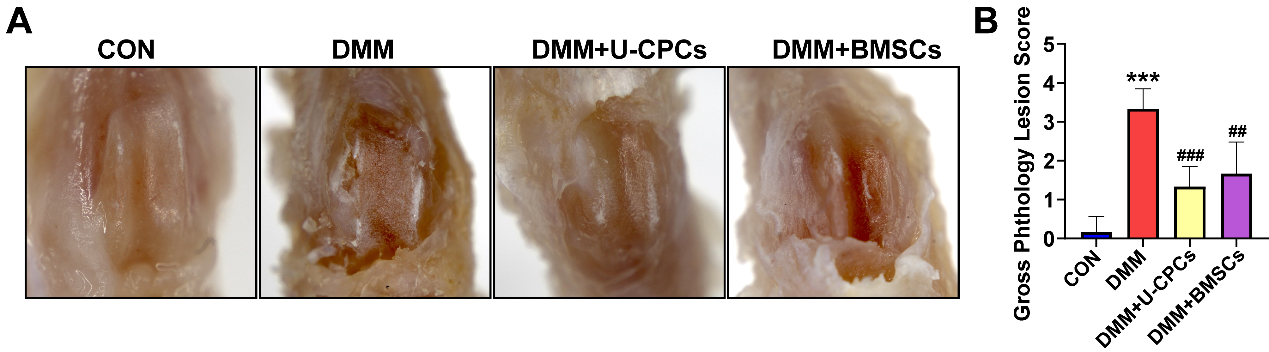

**Figure S11.** Gross assessment of knee joint damage in DMM-induced mice after cell treatment. (A) Representative gross images of knee joints collected 5 weeks after treatment from the control (CON), OA, BMSCs-treated and U-CPCs-treated groups. (B) Quantification of gross joint pathology scores. Data are presented as mean ± SEM; n = 9 per group. ^*^*P*<0.05, ^**^*P*<0.01, ^***^*P*<0.001 *vs* CON; ^#^*P*<0.05, ^##^*P*<0.01, ^###^*P*<0.001 *vs* DMM.

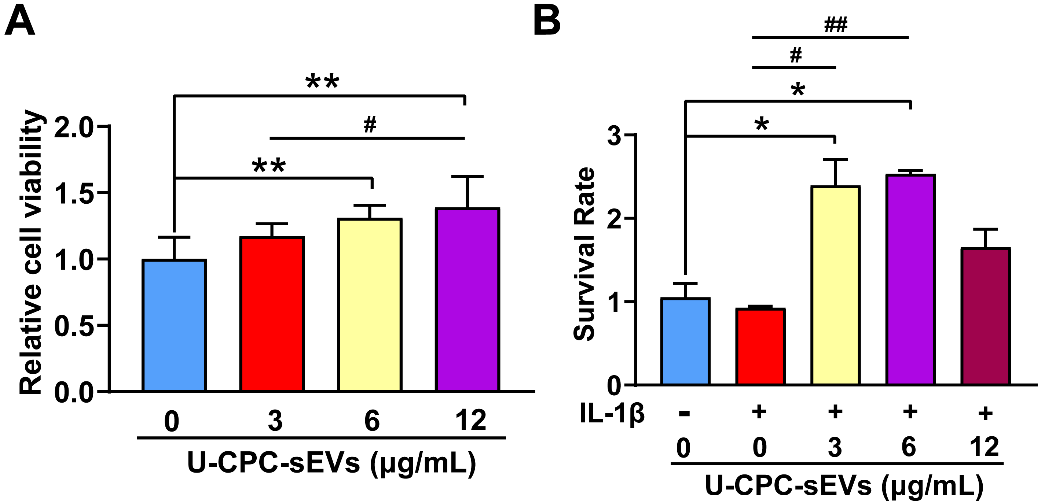

**Figure S12.** Effects of U-CPC-sEVs on C28 Chondrocyte Viability under Basal and Inflammatory Conditions. Cell viability of C28 chondrocytes treated with different concentrations of U-CPC-sEVs in the absence (A) or presence (B) of IL-1β was assessed using the CCK-8 assay. Data are presented as the mean ± SD. ^*^*P* < 0.05, ^**^*P* < 0.01, and ^***^*P* < 0.001; ^#^*P* < 0.05, ^##^*P* < 0.01, and ^###^*P* < 0.001.

**Table S1**. RT-qPCR primer sequences

| Gene | Primer (F=forward; R=reverse; RT=RT Primer Sequence) | | |
| --- | --- | --- | --- |
| ACAN | | | F: ACGGCTTCTGGAGACAGGACTG |
|  |  |  | R: CTGGGATGCTGGTGCTGATGAC |
| COL2A1 | | | F: GGAGCAGCAAGAGCAAGGAGAAG |
|  |  |  | R: TCATCTGGACGTTGGCAGTGTTG |
| SOX9 | | | F: AGGAGAGCGAGGAGGACAAGTTC |
|  |  |  | R: TGTTCTTGCTGGAGCCGTTGAC |
| DUSP7 | | | F: GGCGAGTTCACCTACAAGCAGATC |
|  |  |  | R: GCACCAGGACACCACACTTCTTG |
| DUSP9 | | | F: TGGGCTCCGACTGCTCTGATG |
|  |  |  | R: AGGATCTGGACAGGGAAGGATGC |
| DUSP16 | | | F: CAAGCTGCTGCACCTGGAGAAG |
|  |  |  | R: GCCTTTGTCCTGCTGCCTCTG |
| AKT2 | | | F: GGTCGCCAACAGCCTCAAGC |
|  |  |  | R: ACCGCCACTTCCATCTCCTCAG |
| COL10A1 | | | F: GGACAACAGGGACCCACAGGAG |
|  |  |  | R: TGGACGACCAGGAGCACCATATC |
| MMP13 | | | F: GCGGGAATCCTGAAGGAGAATGC |
|  |  |  | R: ATCGTCAAGTTTGCCAGTCACCTC |
| RUNX2 | | | F: AGGCAGTTCCCAAGCATTTCATCC |
|  |  |  | R: TGGCAGGTAGGTGTGGTAGTGAG |
| Ras | | | F: CACCAGTACAGGGAGCAGATCAAAC  R: GCAGCCAGGTCACACTTGTTCC |
| RafA | | | F: CCTGGCGTTCTGTGACTTCTGC  R: GGCTGTTGGCGGTTGGTACTC |
| RafB | | | F: AAGGGAAAGTGGCATGGTGATGTG  R: TTGAAGGCTTGTAACTGCTGAGGTG |
| Raf1 | | | F: CAAAGAGAGCGGGCACCAGTATC  R: AGAGCCTGACCCAATCCGAGTG |
| MEK1 | | | F: TCATCTGGAGATCAAACCCGCAATC  R: CCATCGCTGTAGAACGCACCATAG |
| MEK2 | | | F: CGCTCACCATCAACCCTACCATC  R: TTCTTCTGCTGCTCGTCAAGTTCC |
| ERK1 | | | F: CATTGTGCAGGACCTGATGGAGAC  R: GTTGGCGGAGTGGATGTACTTGAG |
| c-Fos | | | F: TTCTTCTGCTGCTCGTCAAGTTCC  R: GTCGGTGAGCTGCCAGGATG |
| JNK | | | F: ACTACAGAGCACCCGAGGTCATC  R: TTTCTCCCATGATGCACCCAACTG |
| P38 | | | F: GTGCCCGAGCGTTACCAGAAC  R: GGAGAGCTTCTTCACTGCCACAC |
| U6 | | | F: CTCGCTTCGGCAGCACA |
|  |  |  | R: AACGCTTCACGAATTTGCGT |
| miR-127-3p | | | F: AAGCGGATCGGATCCGTCTGA |
|  |  |  | R: ATCCAGTGCAGGGTCCGAGG |
|  |  |  | RT:GTCGTATCCAGTGCAGGGTCCGAGGTATTCGCACTGGATACGACAGCCAA |
| miR-100-5p | | | F: AACACGTGAACCCGTAGATCCG |
|  |  |  | R: ATCCAGTGCAGGGTCCGAGG |
|  |  |  | RT:GTCGTATCCAGTGCAGGGTCCGAGGTATTCGCACTGGATACGACCACAAG |
| let-7a-5p | | | F: CCACGGAGTGAGGTAGTAGGTTG |
|  |  |  | R: ATCCAGTGCAGGGTCCGAGG |
|  |  |  | RT:GTCGTATCCAGTGCAGGGTCCGAGGTATTCGCACTGGATACGACAACTAT |
| miR-143-3p | | F: AAGCGCCTTGAGATGAAGCACT | |
|  |  | R: ATCCAGTGCAGGGTCCGAGG | |
|  |  | RT:GTCGTATCCAGTGCAGGGTCCGAGGTATTCGCACTGGATACGACGAGCTA | |
| miR-222-3p | | F: AGCGCCTAGCTACATCTGGCT | |
|  |  | R: ATCCAGTGCAGGGTCCGAGG | |
|  |  | RT:GTCGTATCCAGTGCAGGGTCCGAGGTATTCGCACTGGATACGACACCCAG | |

**Table S2**. microRNA sequences

| miRNA | Sequence 1 / Sense (5'-3') | Sequence 2 / Antisense (5'-3') |
| --- | --- | --- |
| miR-127-3p mimic | UCGGAUCCGUCUGAGCUUGGCU | CCAAGCUCAGACGGAUCCGAUU |
| miR-100-5p mimic | AACCCGUAGAUCCGAACUUGUG | CAAGUUCGGAUCUACGGGUUUU |
| let-7a-5p mimic | UGAGGUAGUAGGUUGUAUAGUU | CUAUACAACCUACUACCUCAUU |
| miR-143-3p mimic | UGAGAUGAAGCACUGUAGCUC | GCUACAGUGCUUCAUCUCAUU |
| miR-222-3p mimic | AGCUACAUCUGGCUACUGGGU | CCAGUAGCCAGAUGUAGCUUU |
| DSUP7 siRNA | CGGUCACUGUGGCCUAUCUTT | AGAUAGGCCACAGUGACCGTT |
| Negative control mimic | UUCUCCGAACGUGUCACGUTT | ACGUGACACGUUCGGAGAATT |
| let-7a-5p inhibitor | AACUAUACAACCUACUACCUCA | N/A |
| Negative control inhibitor | CAGUACUUUUGUGUAGUACAA | N/A |
